# D-type cyclins regulate DNA mismatch repair in the G1 and S phases of the cell cycle, maintaining genome stability

**DOI:** 10.1101/2024.01.12.575420

**Authors:** Gergely Rona, Bearach Miwatani-Minter, Qingyue Zhang, Hailey V. Goldberg, Marc A. Kerzhnerman, Jesse B. Howard, Daniele Simoneschi, Ethan Lane, John W. Hobbs, Elizabeth Sassani, Andrew A. Wang, Sarah Keegan, Daniel J. Laverty, Cortt G. Piett, Lorinc S. Pongor, Miranda Li Xu, Joshua Andrade, Anish Thomas, Piotr Sicinski, Manor Askenazi, Beatrix Ueberheide, David Fenyö, Zachary D. Nagel, Michele Pagano

## Abstract

The large majority of oxidative DNA lesions occurring in the G1 phase of the cell cycle are repaired by base excision repair (BER) rather than mismatch repair (MMR) to avoid long resections that can lead to genomic instability and cell death. However, the molecular mechanisms dictating pathway choice between MMR and BER have remained unknown. Here, we show that, during G1, D-type cyclins are recruited to sites of oxidative DNA damage in a PCNA- and p21-dependent manner. D-type cyclins shield p21 from its two ubiquitin ligases CRL1^SKP2^ and CRL4^CDT2^ in a CDK4/6-independent manner. In turn, p21 competes through its PCNA-interacting protein degron with MMR components for their binding to PCNA. This inhibits MMR while not affecting BER. At the G1/S transition, the CRL4^AMBRA1^-dependent degradation of D-type cyclins renders p21 susceptible to proteolysis. These timely degradation events allow the proper binding of MMR proteins to PCNA, enabling the repair of DNA replication errors. Persistent expression of cyclin D1 during S-phase increases the mutational burden and promotes microsatellite instability. Thus, the expression of D-type cyclins inhibits MMR in G1, whereas their degradation is necessary for proper MMR function in S.

**One-Sentence Summary:** To maintain genome stability, D-type cyclins limit mismatch repair (MMR) in G1, whereas their degradation is necessary for proper MMR function in S phase.

## INTRODUCTION

Cyclin-dependent kinases (CDKs) in complex with their regulatory cyclin subunits play a key role in controlling the eukaryotic cell division cycle. D-type cyclins (D1, D2, and D3) and E-type cyclins (E1 and E2) represent the two major subfamilies of mammalian G1 cyclins that, in complex with CDK4/6 and CDK2, respectively, propel the cell through the G1/S transition ^1^. Yet, for reasons that have remained obscure, E-type cyclins persist in S phase to promote DNA replication, whereas the bulk of D-type cyclins is eliminated prior to DNA synthesis ^1–7^.

Apart from cell cycle control, cyclins also regulate homologous recombination repair (HRR) in S and G2 ^8–11^; however, little is known about whether and how cyclins regulate DNA repair in G1. We investigated the role of cyclins in G1 during the response to oxidative stress since this is one of the most frequent causes of DNA damage ^12,13^. BER is the predominant pathway for the repair of oxidative DNA damage ^14^. However, the MMR machinery is also active in G1 (sometimes referred to as noncanonical MMR) ^15–18^ and can also recognize oxidized bases ^19–25^. Thus, MMR and BER share common substrates, but how G1 cells decide which of these pathways will utilize for DNA repair is not known ^26^. Biochemical assays and single molecule studies indicate that MMR involves the excision of DNA tracts whose lengths exceed hundreds of nucleotides ^27,28^. Because of this, as well as the lack of strand discrimination, during G1, repair of oxidized bases by the MMR pathway can result in additional mistakes (*e.g.,* mismatches, reintroduction of oxidized bases, and unligated DNA) ^15,24^. Moreover, it has been proposed that the low availability of nucleotide pools in G1 cells may not allow a complete resynthesis of long-range resected DNA, generating single-stranded DNA tracks that are prone to further damage^15^. Therefore, in the G1 phase of the cell cycle, MMR is an error-prone DNA repair mechanism. Yet, the molecular mechanisms by which MMR is kept in check during G1 and how MMR activity increases in S phase to ensure replication fidelity are unknown.

Here, we present studies that shed light on why mammalian cells eliminate cyclin D1 in S phase and how they favor BER over MMR to repair DNA lesions produced by oxidative stress in G1 in a manner that preserves genome integrity.

## RESULTS

### Cyclin D1 is recruited to DNA lesions in a p21-dependent manner during G1

As a first step to study a possible role of cyclins in DNA repair during G1, we tested whether the major human cyclins (A2, B1, D1, D2, D3, E1, and F, which are encoded by *CCNA2, CCNB1, CCND1, CCND2, CCND3, CCNE1,* and *CCNF*, respectively) are recruited to laser-induced DNA damage sites. Laser micro-irradiation was used as it induces at the same time a variety of lesions, including single-stranded breaks (SSBs), base oxidation, UV adducts, double-stranded breaks (DSBs) *etc*. During live-cell imaging, we initially classified interphase cells into S phase cells (positive for PCNA foci) and non-S phase cells (*i.e.,* G1 and G2 cells that are negative for PCNA foci). Cyclin D1 and, to a lesser extent, cyclin D2 and cyclin D3, were recruited to DNA lesions in cells without PCNA foci, but not in S phase cells (Fig. 1A and Fig. S1A-B). Similar observations were made using confocal microscopy (Fig. S1C). In contrast, none of the other cyclins were recruited in any phase of the cell cycle (Fig. 1A and Fig. S1A). The nuclear translocation of cyclin D1 and its interaction with PCNA depend on p21, a cyclin interactor belonging to the KIP (kinase inhibitory protein) family ^29–31^. Depletion of p21, but not its paralogs p27 and p57 inhibited the recruitment of cyclin D1 to laser-induced damage sites (Fig. 1B and Fig. S1D-E). Silencing of p21 also negatively affected the nuclear accumulation of cyclin D1; however, even when fused to a nuclear-localization signal (NLS) cyclin D1 was unable to be recruited to DNA lesions, despite accumulating in the nucleus independently of p21 (Fig. 1B). Similar observations were made using parental (*p21*^+/+^) and *p21*^-/-^ cells (Fig. 1C and Fig. S1F). A mutant of cyclin D1(HP) that does not bind to p21 and, therefore, PCNA ^31^ was unable to localize to laser-induced DNA damage sites even when forced into the nucleus by an additional NLS (Fig. 1D and Fig. S1G-I). Cyclin D2 and cyclin D3 had a similar p21-dependent recruitment pattern (Fig. 1C).

**Figure 1.**
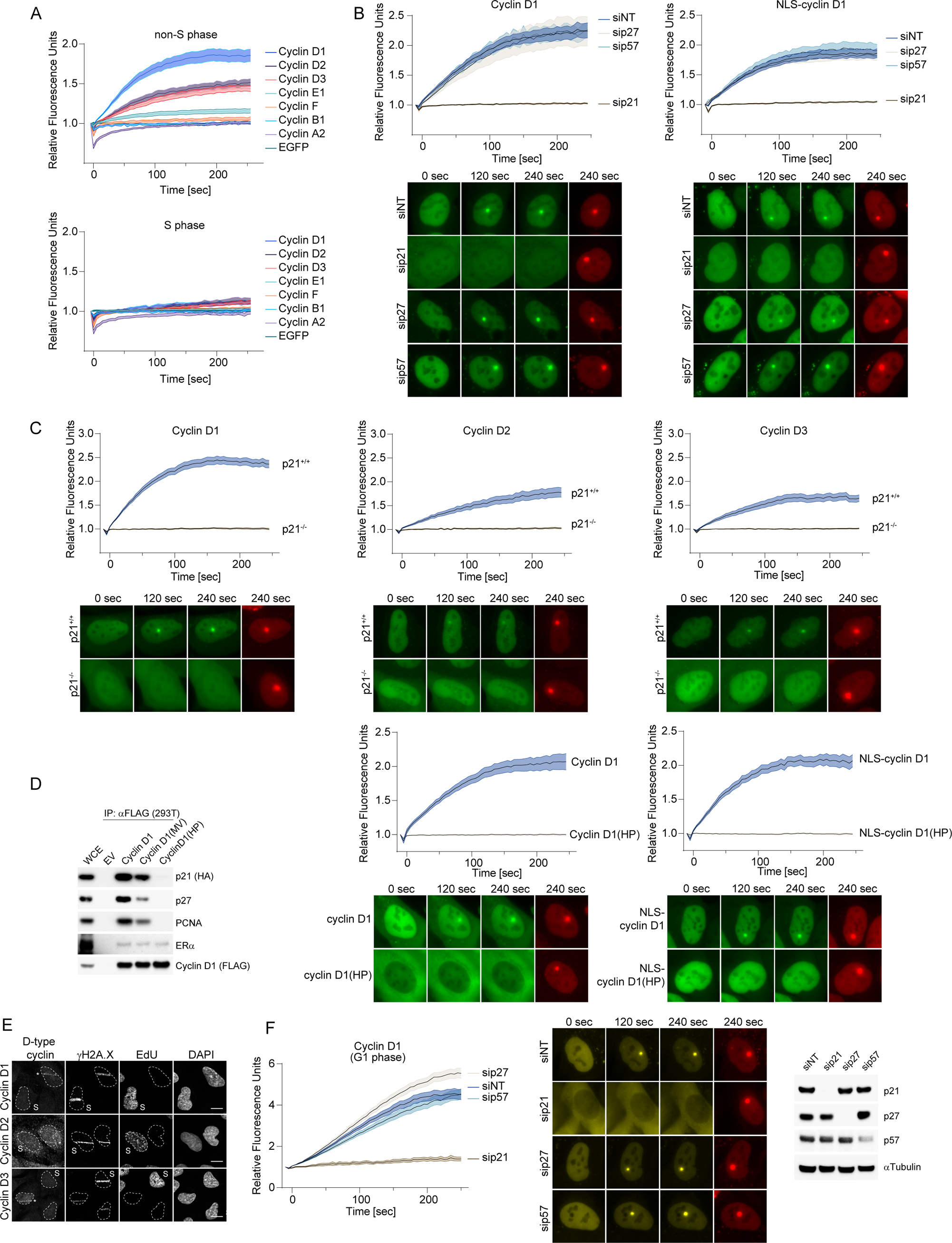
D-type cyclins are recruited to DNA lesions in a p21 dependent manner. A) U-2OS cells stably expressing EGFP-tagged cyclins, EGFP, and mPlum-PCNA were pre-sensitized with BrdU for 24 hours and subjected to 405 nm laser induced damage. DNA damage recruitment dynamics were captured by live cell imaging. Non-S phase cells (upper panel) and S phase cells (bottom panel) were identified based on the presence of PCNA foci. Relative fluorescence values and images were acquired every 5 seconds for 4 minutes. For each condition, ≥25 cells were evaluated from 3 independent experiments. Mean relative fluorescence values and standard errors were plotted against time. Representative images are shown in Fig. S1A. Times are indicated in seconds. B) U-2OS cells stably expressing either mAzGreen-cyclin D1 or NLS-mAzGreen-cyclin D1 (in green) together with mPlum-PCNA (in red) were transfected with siRNAs targeting p21, p27, p57 or a non-targeting control (NT). Cells were treated and analyzed as in *(A)*. Representative images are shown below the graphs. C) Parental (*p21*^+/+^) or *p21^-/-^* U-2OS cells stably expressing mAzGreen-cyclin D1, D2, or D3 and mPlum-PCNA were treated and analyzed as in *(A)*. D) Left: HEK 293T cells were co-transfected with the indicated FFSS-cyclin D1 constructs together with either p21-HA or EV. Cyclin D1(MV) denotes mutations M56A and V60A. Cyclin D1(HP) denotes mutations M56A, V60A, and W63A. Cell lysates were immunoprecipitated with an anti-FLAG resin, followed by elution using 3× FLAG peptide. Immunoprecipitates were probed with the indicated antibodies. Whole cell extract (WCE) is of the EV sample. Middle and right panels: U-2OS cells stably expressing the indicated mAzGreen-cyclin D1 constructs and mPlum-PCNA were treated and analyzed as in *(A)*. E) Confocal images of U-2OS cells fixed 2 minutes after laser micro-irradiation and stained for either the DNA damage marker γH2A.X, the replication marker EdU and the indicated D-type cyclins. DAPI was used to counterstain nuclear DNA. A white dashed line denotes the border of each nucleus. Scale bar represents 5 μm. F) G1-synchronized RPE1 cells harboring fluorescent-tagged endogenous proteins (mRuby-PCNA, mVenus-cyclin D1 and p21-mTurquoise2) were treated and analyzed as in *(A)*. Representative images are shown in the middle panel. The efficiency of depletion is shown in the right panel.

Next, we synchronized cells in G1 and observed that cyclin D1 was also recruited to laser-induced DNA damage sites in a p21-dependent manner, while cyclin E1 was not recruited under identical conditions (Fig. S1J). Recruitment of cyclin D1 in G1 was independent of CDK4/6 activity, since it was not affected by treatment of cells with a CDK4/6 inhibitor, palbociclib (Fig. S1K).

Using immunofluorescence, we confirmed that endogenous cyclin D1, D2, and D3 also localized to laser-induced DNA lesions in non-S phase cells (as indicated by their EdU-negative nuclei) (Fig. 1E). In synchronized G1 cells, endogenous cyclin D1 was also recruited in a p21-dependent manner to DNA-lesions (Fig. 1F). Using a PIP (PCNA-interacting protein) box mutant of p21(ΔPCNA), we showed that p21’s recruitment depends on its ability to bind PCNA (Fig. S1L-N), in agreement with previous reports ^32^. An RxL mutant of p21(RxL), which binds to PCNA but not to cyclins, was still able to localize to laser-induced DNA lesions (Fig. S1L-N), showing that its ability to bind cyclins is not needed for recruitment.

These observations suggest a hierarchical recruitment model of PCNA à p21 à D-type cyclins upon induction of DNA damage in G1.

### D-type cyclins are recruited to oxidative DNA lesions in the proximity of BER components

Laser-induced microirradiation produces a variety of DNA lesions. To determine the type(s) of DNA lesion to which cyclin D1 is recruited, we treated G1-synchronized cells with various DNA damaging agents before fractionating cell lysates into soluble and chromatin bound fractions (Fig. 2A and Fig S2A). Cyclin D1, D2 and D3 were detected in the chromatin fraction upon oxidative stress (H_2_O_2_). They also localized to chromatin to a lesser extent upon treatment with an alkylating DNA damaging agent (MMS), but were not recruited with bleomycin and NCS, which induce DSBs and SSBs (Fig. 2A and Fig S2A). Upon oxidative stress, chromatin recruitment of both endogenous or exogenous D-type cyclins was largely dependent on p21 (Fig. 2B and Fig. S2B). The residual recruitment of D-type cyclins in *CDKN1A*^-/-^ cells (*CDKN1A* encodes p21) was almost completely abolished by silencing p57 (Fig. S2C), which contains a PIP box and seemed to partially substitute for p21 in knockout cells.

**Figure 2.**
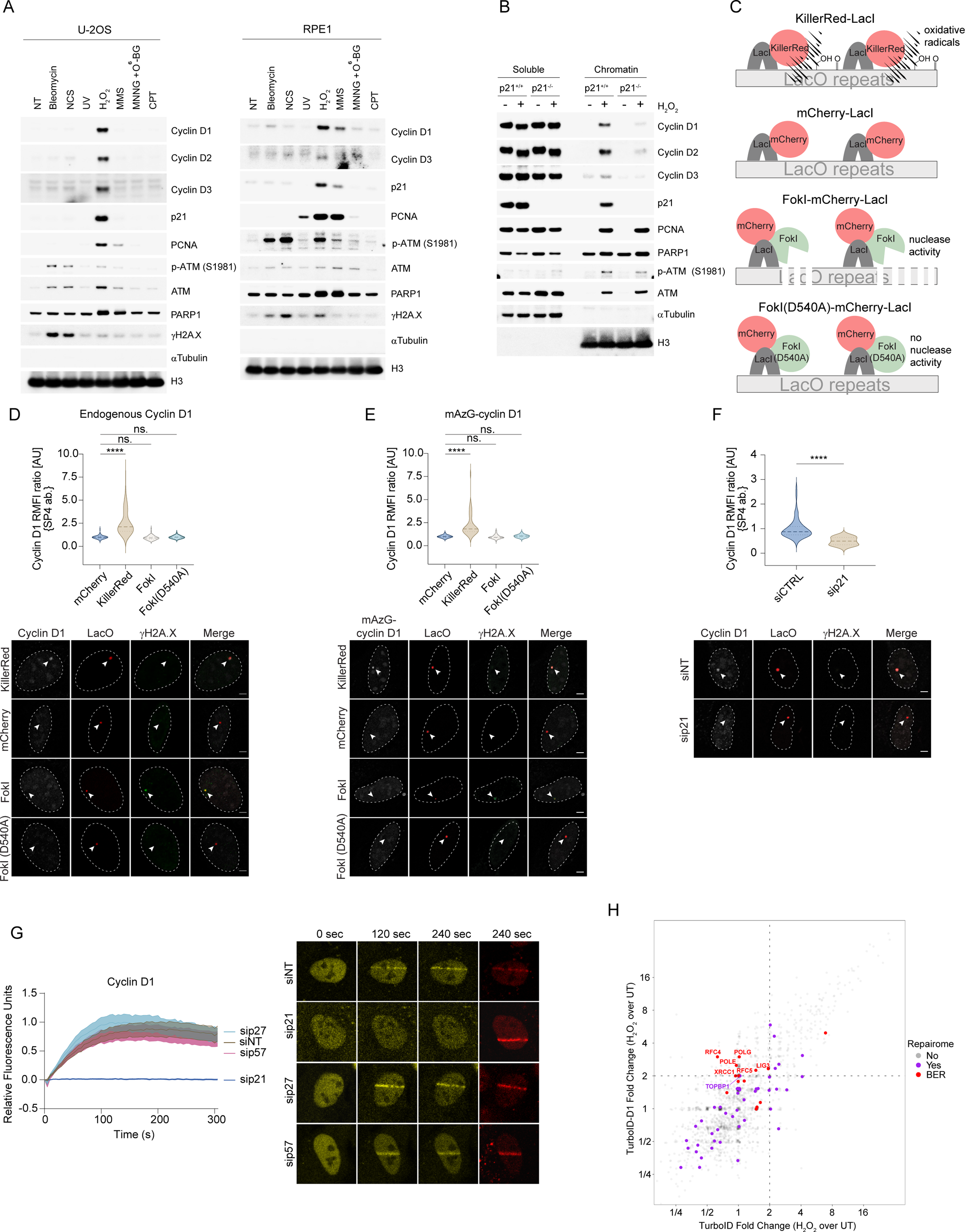
D-type cyclins are recruited to oxidative DNA lesions in the proximity of BER components. A) G1-synchronized U-2OS and RPE1 cells were treated with the indicated DNA damaging agents. Cells were then fractionated into soluble and chromatin fractions, and lysates were immunoblotted as indicated. The figure shows the chromatin fractions while Fig. S2A shows the soluble fractions. B) G1-synchronized parental (*p21*^+/+^) and *p21^-/-^* U-2OS cells were treated with 100 μM H_2_O_2_ for 20 minutes. Cells were then fractionated into soluble and chromatin fractions, and immunoblotted as indicated. C) Schematic of the site-specific DNA damage recruitment assay. 2-6-3 U-2OS reporter cells containing lac operator (LacO) repeats integrated into their genome were transfected with vectors expressing either mKillerRed-LacI or mCherry-LacI-FokI to generate local DNA oxidization and DSBs, respectively. mCherry-LacI and a nuclease FokI(D540A) inactive mutant were used as negative controls. D) 2-6-3 U-2OS cells were transfected with the indicated constructs and immunostained for γH2A.X (green) and endogenous cyclin D1 (grey). The latter was detected using the SP4 rabbit monoclonal antibody. White arrowheads indicate the position of the LacO (red) in the nucleus, which is outlined by a white dashed line. Quantification of the cyclin D1 relative mean fluorescence intensity (RMFI) was carried out from 3 independent experiments as in ^83^ and visualized on violin plots. Dashed lines represent the median on the plots. Scale bar represents 2 μm. E) 2-6-3 U-2OS cells were co-transfected with mAzGreen-cyclin D1 (grey) and the indicated constructs, and subsequently immunostained for γH2A.X (green). White arrowheads indicate the position of the LacO (red) in the nucleus, which is outlined by white dashed line. Quantification of the mAzGreen-cyclin D1 relative mean fluorescence intensity (RMFI) was carried out from 3 independent experiments and visualized on violin plots. Dashed lines represent the median on the plots. Scale bar represents 2 μm. F) 2-6-3 U-2OS cells were transfected with the indicated constructs and siRNAs targeting p21 or a non-targeting control (NT), and then immunostained for γH2A.X (green) and cyclin D1 (grey). The latter was detected and quantified as in *(D).* G) RPE1 cells harboring fluorescent-tagged endogenous proteins (mRuby-PCNA, mVenus-cyclin D1, and p21-mTurquoise2) were micro-irradiated as described in ^34^. DNA damage recruitment dynamics were captured by live-cell imaging. Non-S phase cells were identified based on the absence of PCNA foci and analyzed as in Fig. 1A. H) Scatter plot of fold-changes in PSM (peptide spectral match) counts of H_2_O_2_-treated vs. non-treated cells for the control TurboID or TurboID fused to cyclin D1 expressed in U-2OS cells. Raw values are reported in Supplementary Table S1. Known DNA repair proteins are highlighted in purple, except BER proteins that are in red. Selected proteins must have a fold-change greater than or equal to 2 in the case of cyclin D1 and not have a fold-change greater than or equal to 2 in the case for TurboID on its own. Within these selected proteins, only the BER pathway was significantly enriched (p-value = 0.005711, Fisher’s exact test) among the DNA repair pathways (BER, NER, MMR, HRR, NHEJ, FAN DR, TLS, and DDA).

To further verify the recruitment of cyclin D1 to oxidative DNA lesions, we used LacI-fused KillerRed to induce local accumulation of oxidized bases in a LacO array in the genome of 2-6-3 U-2OS reporter cells ^33^ (Fig. 2C). LacI-KillerRed was able to recruit OGG1 but did not induce a γH2A.X signal (indicative of DSBs), highlighting its specificity for generating oxidized bases (Fig. S2D-E). LacI-KillerRed was able to recruit cyclin D1 in a p21-dependent manner, whereas LacI-FokI (inducing DSBs), a LacI-FokI(D540A) inactive nuclease mutant, or LacI-Cherry (another negative control) did not induce the recruitment of cyclin D1 (Fig. 2D-F and Fig. S2F-G). Finally, when micro-irradiation was used under specific conditions that do not induce DSBs ^34^, endogenous cyclin D1 was still recruited to DNA damage sites (Fig. 2G).

To understand which DNA repair pathway is active at the lesions where cyclin D1 recruits upon H_2_O_2_ treatment, proximity biotinylation assays ^35,36^ were carried out by fusing the enzyme BirA to cyclin D1. Proteins in the proximity of chromatin-bound cyclin D1 were labeled by biotinylation, purified, and analyzed by mass-spectrometry. Among the DNA damage and repair pathways, only the BER pathway was significantly enriched in the proximity of chromatin-bound cyclin D1 upon H_2_O_2_ treatment (Fig. 2H and Fig. S2H-I).

These observations suggest that, upon oxidative stress, D-type cyclins accumulate on the chromatin in the proximity of members of the BER pathway.

### Cyclin D1 negatively regulates the binding of MMR proteins to PCNA

For more than 30 years it has been known that PCNA and cyclin D1 are present in the same protein complex ^37^, but the significance of this interaction is not clear ^38^. PCNA acts as a scaffolding protein for components of several DNA repair pathways ^39^. Since the recruitment of D-type cyclins upon oxidative DNA damage is p21- and PCNA-dependent (Fig. 1,2), we sought to determine the possible changes in the DNA damage-induced PCNA interactome in the presence or absence of D-type cyclins in G1. We found that, upon depletion of D-type cyclins, the only DNA repair proteins that were increased in the proximity of PCNA were MMR proteins, with MSH2, MSH3, and MSH6 increasing 2.3, 3.9, and 2.2 folds, respectively, compared to control samples (Fig. 3A and Fig. S3A-B). We validated these findings using immunoprecipitation followed by immunoblotting in cells that were synchronized in G1. Endogenous PCNA was immunoprecipitated from the chromatin bound fraction with or without H_2_O_2_ treatment, and probed for MMR interactors (Fig. 3B,C and Fig. S3C,D). When cells were depleted of D-type cyclins, MMR components co-immunoprecipitated more abundantly with PCNA (Fig. 3B and Fig. S3C; compare lanes 3 and 4). When G1 cells were depleted of p21, p27, and p57, we observed similar changes in binding between PCNA and MMR components (Fig. 3B and Fig. S3C; compare lanes 3 and 5). We then used cells lacking AMBRA1, the ubiquitin ligase targeting D-type cyclins for degradation, that have high levels of endogenous D-type cyclins and p21 ^40–42^. In these cells, we observed the opposite trend, whereby PCNA bound less MMR components in G1 (Fig. 3C and Fig. S3D; compare lanes 3 to 4 and 5).

**Figure 3.**
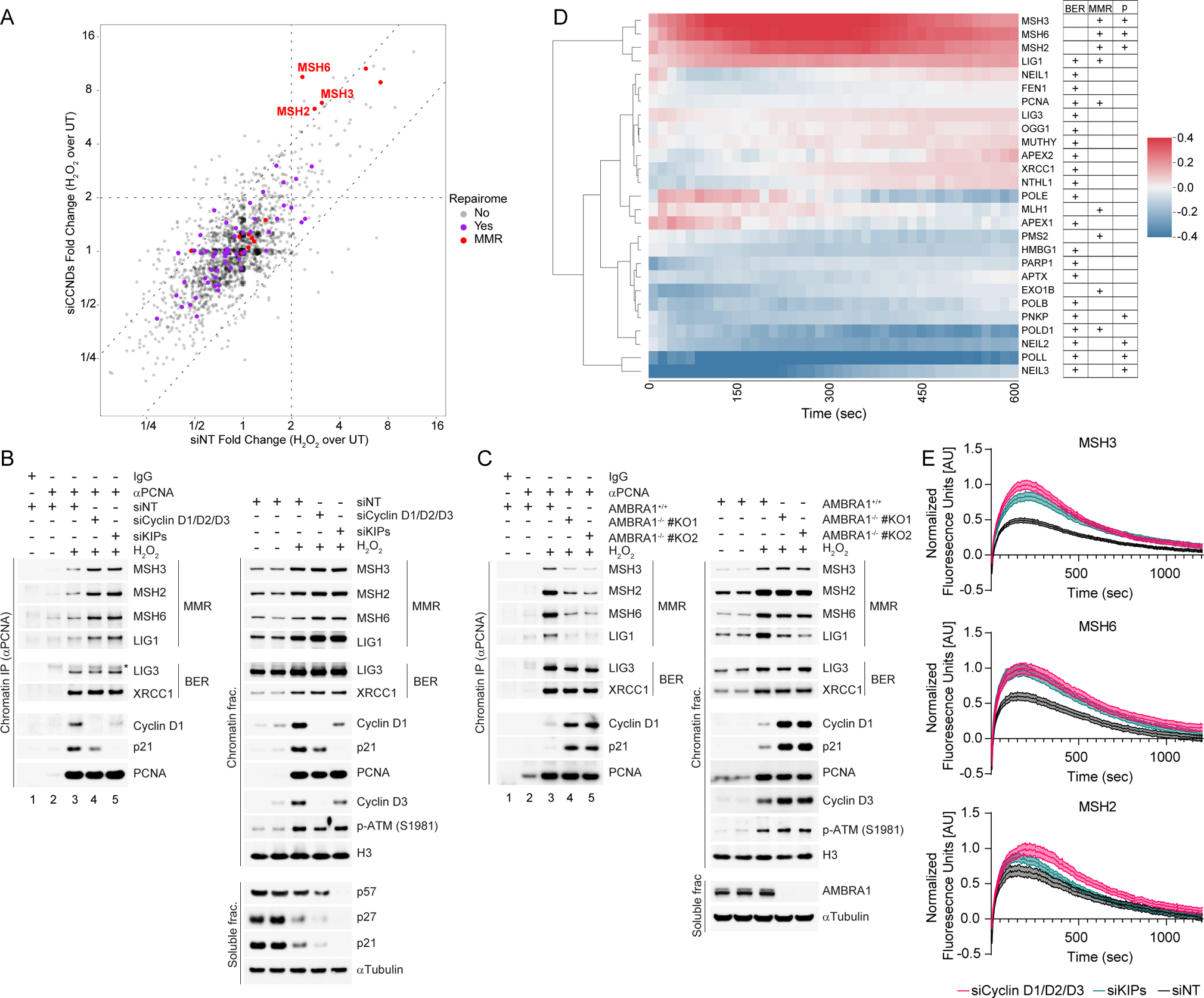
D-type cyclins inhibit the binding of MMR proteins to PCNA. A) Scatter plot of fold-changes in PSM (peptide spectral match) counts of H_2_O_2_-treated vs. non-treated cells of TurboID-PCNA expressed in RPE1 cells. Cells were transfected with either non-targeting (siNT) or siRNA targeting D-type cyclins (siCCNDs) and subsequently synchronized in G1. Selected proteins must have a fold-change greater than two-fold in both cases, but to be considered enriched upon siCCNDs, the fold-changes must be at least twice as high in the case of siCCNDs compared to siNT. Raw values are reported in Supplementary Table S2. B) RPE1 cells were transfected with either non-targeting (siNT) or siRNA targeting D-type cyclins or KIPs, and subsequently synchronized in G1. Cells were then treated with 100 μM H_2_O_2_ before fractionated into soluble and chromatin fractions. PCNA was immunoprecipitated from the chromatin fraction and co-purified proteins were immunoblotted as indicated (left panel). Right panels show chromatin and soluble fractions. Asterix denotes a non-specific band. C) G1-synchronized parental (*AMBRA1*^+/+^) or *AMBRA1*^-/-^ RPE1 cells were treated with 100 μM H_2_O_2_ before fractionated into soluble and chromatin fractions. PCNA was immunoprecipitated from the chromatin fraction and co-purified proteins were immunoblotted as indicated (left panel). Right panels show chromatin and soluble fractions. D) U-2OS cells stably expressing mCherry-tagged BER pathway components or AcGFP-tagged MMR pathway components were transfected with either non-targeting or siRNA targeting D-type cyclins, and subsequently synchronized in G1. Each row in the heatmap shows recruitment of a BER or MMR protein at micro-irradiation sites that is higher (red) or lower (blue) in D-type cyclin depleted cells when compared to control cells. Columns represent standardized recruitment values that were averaged over 15 second intervals. Column label units refer to seconds post-irradiation. Hierarchical clustering of rows in the heatmap was performed with Euclidean distance measure and Ward clustering method. The table on the right shows if the protein of interest is part of the BER or MMR pathway. The “p” column indicates if the mean intensity of the normalized difference between control and D-type cyclin-depleted cells was significant over a window of 60 seconds across the entire 10 minutes (+: p < 0.01). E) U-2OS cells stably expressing AcGFP-tagged MSH2, MSH3, and MSH6 were transfected with siRNAs targeting D-type cyclins, KIPs or a non-targeting control (NT), and subsequently synchronized in G1. DNA damage recruitment dynamics were captured by live-cell imaging. Relative fluorescence values and images were acquired every 5 seconds for 10 minutes. Mean relative fluorescence values and standard errors were plotted against time. Times are indicated in seconds.

We also used live-cell imaging and laser micro-irradiation to establish the consequences of altered D-type cyclins levels more accurately on the recruitment dynamics of MMR and BER components to DNA lesions in G1. MMR and BER components were tagged with fluorescent proteins, and their recruitment dynamics were evaluated with and without the silencing of D-type cyclins. In agreement with our earlier data, when depleting D-type cyclins, there was a significant increase in the recruitment of MSH2, MSH3, and MSH6 (Fig. 3D-E and Fig. S3E). We also observed a significant upregulation in the recruitment of MSH2, MSH3 and MSH6, when cells were depleted of KIPs (Fig. 3E). The recruitment of some BER factors (PNKP, POLL, NEIL2 and NEIL3), but not BER core components (XRCC1, POLB, and LIG3), were significantly reduced (Fig. 3D and Fig. S3E). Similarly, the binding of PCNA to BER core components was not affected by altering the levels of D-type cyclins or KIPs (Fig. 3B,C and Fig. S3D,E).

These observations suggest that upon oxidative stress, D-type cyclins inhibit the interaction between PCNA and MMR proteins while not affecting the core BER components.

### D-type cyclins promote the repair of oxidative DNA lesions during G1

Since D-type cyclins are recruited to oxidative lesions, we ought to investigate their role in oxidative DNA damage repair. Using a comet assay, we observed that while control G1 cells repaired most of their oxidative stress-induced DNA lesions, G1 cells with reduced levels of either D-type cyclins or KIPs were not able to repair their DNA to a similar extent (Fig. 4A-B and Fig. S4A) and displayed apoptotic markers (Fig. S4B).

**Figure 4.**
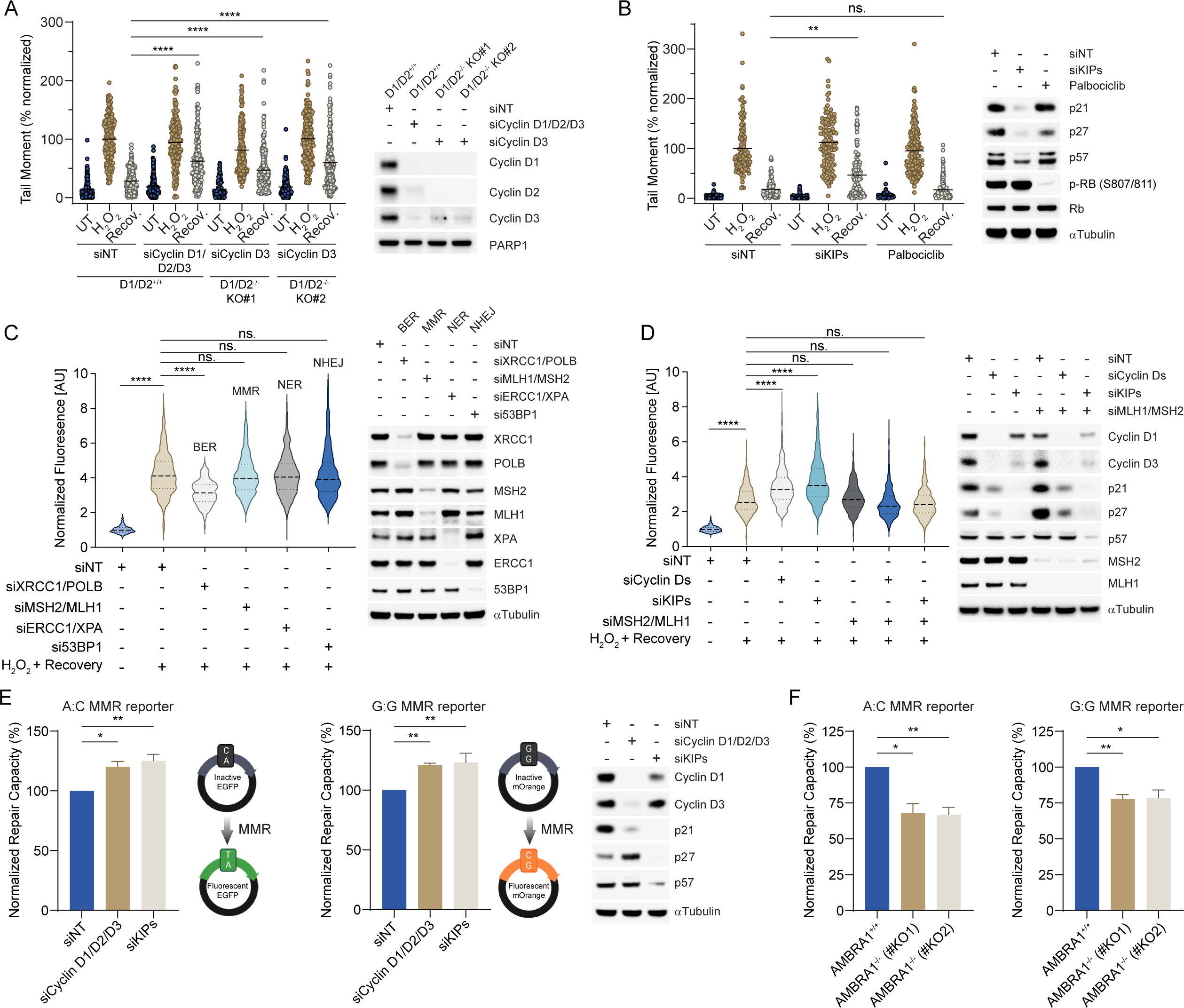
D-type cyclin levels promote the repair of oxidative DNA lesions during G1. A) Two *CCND1^-/-^; CCND2^-/-^* (*D1/D2^-/-^*) U-2OS clones (clones #KO1 and #KO2) and parental U-2OS cells (*D1/D2^+/+^*) were transfected with siRNAs targeting D-type cyclins, cyclin D3, or a non-targeting control (siNT), and subsequently synchronized in G1. Cells were then treated with H_2_O_2_ for 15 minutes or left untreated (UT). Half of the samples treated with H_2_O_2_ were allowed to recover in media without H_2_O_2_ for 4 hours (Recov.), after which all samples were subjected to comet assay. Comet tail moment values for were normalized to the siNT “H_2_O_2_” samples. Lines represent mean values on the plots from 4 independent experiments. The efficiency of depletion for cyclin D1, cyclin D2, and cyclin D3 is shown on the right panel. B) U-2OS cells were transfected with siRNAs targeting KIPs, or a non-targeting control (siNT), and subsequently synchronized in G1. Cells were then treated with H_2_O_2_ for 15 minutes or left untreated (UT). Half of the samples treated with H_2_O_2_ were allowed to recover in media without H_2_O_2_ for 4 hours (Recov.), after which all samples were subjected to comet assay. Where indicated, cells were pretreated for 2 hours with palbociclib (CDK4/6i) and kept in palbociclib during the 4 hours recovery. Comet tail moment values were normalized to the siNT “H_2_O_2_” samples. Lines represent mean values on the plots from 3 independent experiments. The efficiency of p21, p27, p57 depletion and CDK4/6 inhibition is shown on the right panel. C) RPE1 cells were transfected with either siRNAs targeting the indicated proteins or a non-targeting siRNA control (siNT), and subsequently synchronized in G1. Where indicated, cells were treated with H_2_O_2_ for 30 minutes and then allowed to recover for 1.5 hours in DMEM supplemented with EdU. The EdU signal was normalized to the untreated siNT samples. Measurements were carried out from 3 independent experiments and visualized on violin plots. Dashed lines represent the median on the plots. The efficiency of the siRNA knockdowns is shown using immunoblotting on the right. D) RPE1 cells were transfected with either siRNAs targeting D-type cyclins, KIPs, MLH1, MSH2, or a non-targeting siRNA control (siNT), and subsequently synchronized in G1. Cells were then treated and evaluated as in *(C).* E) RPE1 cells were transfected with siRNAs targeting D-type cyclins, KIPs, or a non-targeting control (siNT) and serum-starved to arrest them in G0. Cells were then nucleoporated with the indicated FM-HCR reporter plasmids and released into serum containing media. Ten hours later, cells, now in G1, were subjected to FACS analysis. DNA repair activity for each siRNA was normalized to siNT and was represented as normalized repair activity. Graphs show average and standard deviation from at least 3 independent experiments. Schematic of the transcriptional mutagenesis-based fluorescent reporters is shown next to the graphs. The base pairs shown correspond to sites that code for a key amino acid of the chromophores of the fluorescent proteins. The transcribed strand is on the top. The efficiency of the siRNA knock-downs is shown in the right panel. F) Parental (*AMBRA1*^+/+^) and *AMBRA1*^-/-^ RPE1 cells (clones #KO1 and #KO2) were treated and analyzed as in *(E)*.

We also measured DNA repair synthesis by EdU incorporation and found that in response to oxidative stress in G1 cells, depletion of BER proteins, but not proteins involved in MMR and other repair pathways, inhibited DNA repair synthesis (Fig. 4C). This result confirms previous reports showing that the BER pathway is the predominant repair pathway acting on oxidized bases in G1 ^14,17^, despite the MMR pathway is also active in this phase of the cell cycle ^15–18^. Depleting D-type cyclins or KIPs led to an increase in DNA repair synthesis compared to control cells (Fig. 4D). Importantly, this increase in EdU incorporation was driven by MMR, since the depletion of MSH2 and MLH1 rescued this phenotype (Fig. 4D). Finally, inhibition of DNA repair synthesis by depletion of BER proteins was rescued by the co-silencing of D-type cyclins (Fig. S4C). A possible explanation for this result is that the removal of D-type cyclins allows MMR proteins to act on these lesions.

To directly measure the effect of D-type cyclin levels on MMR activity, we used the fluorescence-based multiplex host cell reactivation (FM-HCR) assay to measure MMR activity in live cells ^43,44^. The assay is highly validated in multiple cell lines and can detect modest alterations in mismatch repair capacity ^45,46^. In brief, fluorescent reporter plasmid substrates, each containing a single, site-specific DNA lesion, were transfected into cells whereupon DNA repair alters the fluorescent reporter signal, which is detected by flow cytometry (Fig. S4D). Consistent with previous validation ^44^, the FM-HCR assay showed ∼50% reduction in MMR activity in cells depleted of both MSH2 and MLH1, and a ∼90-95% reduction in activity in *MLH1*^-/-^ cells (Fig. S4E-F). Notably, the closed circular FM-HCR reporter plasmids confirm MMR-dependent processing of a single base mispair in the absence of replication or a strand discrimination signal. Depletion of D-type cyclins or KIPs in G1 cells significantly increased MMR activity of both a G:G mismatch and an A:C mismatch when compared to non-targeting siRNA transfected cells (Fig. 4E). In contrast, the *AMBRA1*^-/-^ cells displayed a significant reduction in MMR activity (Fig. 4F). Accordingly, when overexpressing either D-type cyclins or p21, but not p21(ΔPCNA), we observed a reduction in the MMR activity (Fig. S4G). Depletion of p21 in *AMBRA1*^-/-^ cells rescued the reduction in the MMR capacity (Fig. S4H). Note that similar moderate alterations in mismatch repair capacity are sufficient to induce major effects on cellular sensitivity to both mutagenesis and killing with alkylating agents ^45,46^.

These observations suggest that D-type cyclins inhibit the cell’s MMR activity in G1 and contribute to the proper DNA repair and recovery after oxidative genotoxic stress.

### D-type cyclins stabilize p21, which competes with MMR proteins to bind PCNA

To understand the mechanism by which D-type cyclins regulate MMR, we first investigated if the kinase activity of their associated catalytic subunits (CDK4 and CDK6) were involved in any of the observed phenotypes. The binding between PCNA and MMR components (detected by immunoprecipitation/immunoblot), the overall DNA repair capacity (as measured by comet assay), DNA synthesis repair (measured by EdU incorporation), and MMR activity (measured by FM-HCR) were not altered by inhibiting CDK4/6 with palbociclib or depleting these kinases with siRNA in G1 synchronized cells (Fig. 4B and Fig. S5A-C). Therefore, the role of D-type cyclins in the repair of oxidative lesions appears to be independent of the associated CDK activity.

We noticed that when depleting D-type cyclins, we also reduced the levels of p21 (Fig. 3B, Fig. S3C, Fig. 4D, Fig. 4E, Fig. S4B and Fig. S4C). Accordingly, *AMBRA1*^-/-^ cells, which have high levels of D-type cyclins, display increased p21 levels (Fig. 3C, Fig. S3D, Fig. S4D, Fig. S5C) similarly to when D-type cyclins are overexpressed (Fig. S4G). The observation that p21 mRNA levels did not change upon depletion of D-type cyclins (Fig. S5D) suggests a non-transcriptional regulation of p21 levels by AMBRA1 and D-type cyclins. Indeed, compared to parental cells, *AMBRA1*^-/-^ cells displayed more stable p21, as determined by inhibiting translation with cycloheximide (Fig. 5A and Fig. S5E). This effect could be largely rescued by silencing D-type cyclins in *AMBRA1*^-/-^ cells (Fig. 5B and Fig. S5F). Moreover, silencing D-type cyclins destabilized p21 in G1 synchronized cells (Fig. 5C and Fig. S5G). We observed a similar destabilization of p21 in *CCND1*/*2*/*3* triple knockout MEFs as compared to parental MEFs (Fig. S5H).

**Figure 5.**
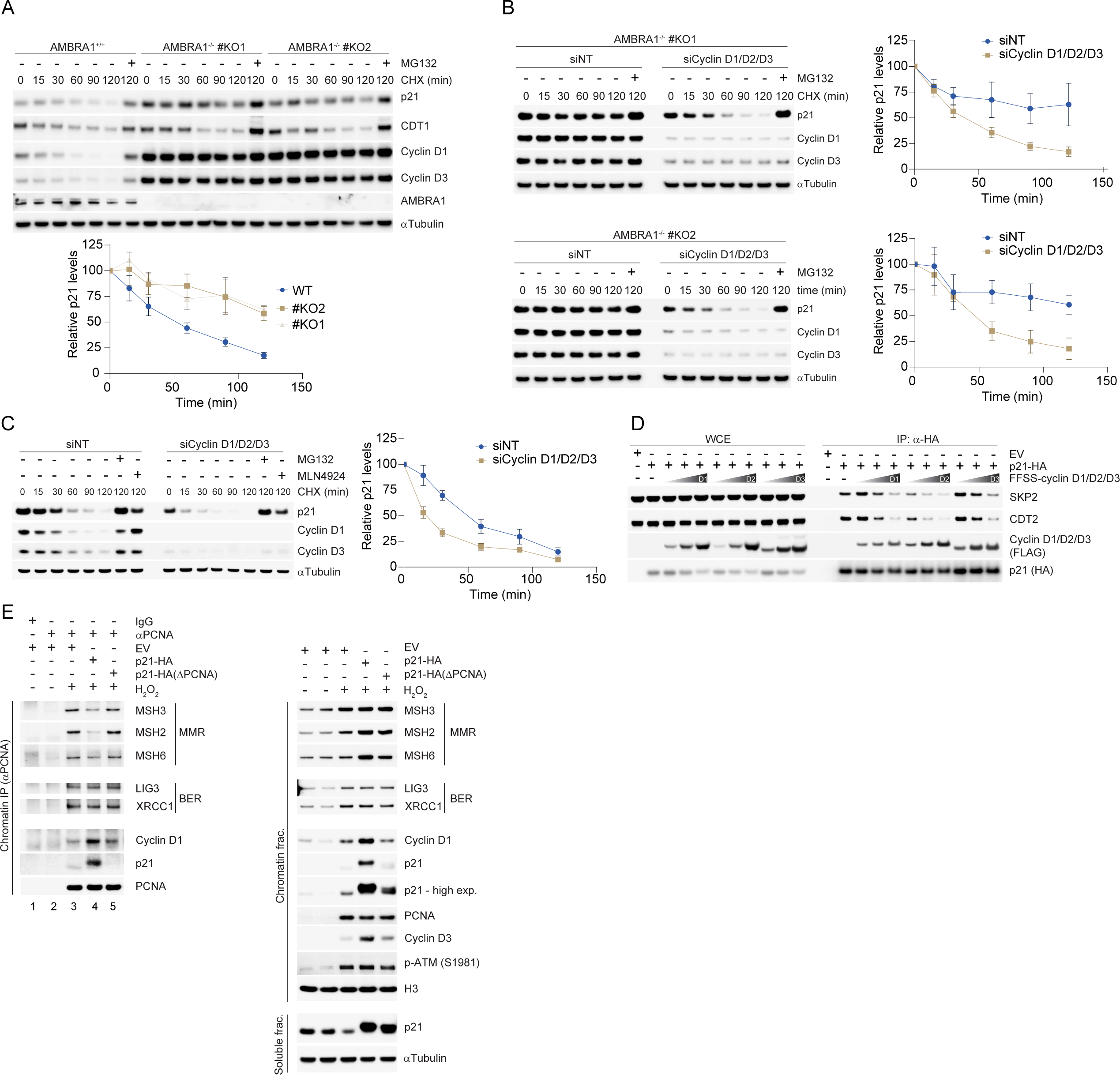
D-type cyclins stabilize p21, which competes with MMR proteins to bind PCNA. A) Parental (*AMBRA1*^+/+^) and *AMBRA1*^-/-^ RPE1 cells (clones #KO1 and #KO2) were treated with cycloheximide (CHX) and MG132 as indicated, and lysates were blotted with the indicated antibodies. The graph at the bottom shows the quantification of p21 levels from three independent experiments. Error bars indicate standard error. B) *AMBRA1*^-/-^ RPE1 cells (clones #KO1 and #KO2) were transfected with siRNAs targeting D-type cyclins, or a non-targeting control (siNT) before being treated with cycloheximide (CHX) and MG132 as indicated. Lysates were blotted with the indicated antibodies. The graphs on the right show the quantification of p21 levels from three independent experiments. Error bars indicate standard error. C) RPE1 cells were transfected with siRNAs targeting D-type cyclins or a non-targeting control (siNT), and subsequently synchronized in G1. Cells were then treated with cycloheximide (CHX) and either MG132 or MLN4924 as indicated. Lysates were blotted with the indicated antibodies. The graph on the right shows the quantification of p21 levels from three independent experiments. Error bars indicate standard error. D) HEK 293T cells were transfected with either an empty vector (EV) or HA-p21, together with increasing amounts of FFSS-cyclin D1, FFSS-cyclin D2, or FFSS-cyclin D3 vectors. p21 was immunoprecipitated from the lysates using HA-beads and co-purified proteins were immunoblotted as indicated. E) Stably transduced RPE1 cells harboring a doxycycline-inducible HA-p21, HA-p21(ΔPCNA), or EV were synchronized into G0 using serum starvation, during which the transgenes were induced by doxycycline. After serum release, G1 phase cells were treated with 100 μM H_2_O_2_ before fractionating them into soluble and chromatin fractions. PCNA was immunoprecipitated from the chromatin fraction and co-purified proteins were immunoblotted as indicated (left panel). Right panels show chromatin and soluble fractions.

We hypothesized that D-type cyclins can shield p21 from its two cullin-RING ubiquitin ligases, CRL1^SKP2^ and CRL4^CDT2^ ^47,48^, and therefore stabilize it. When immunoprecipitating p21 in the presence of increasing doses of either cyclin D1, D2 or D3, we observed a reduced binding of p21 to SKP2 and CDT2 (Fig. 5D), supporting our hypothesis.

Increasing the levels of D-type cyclins not only resulted in the stabilization of p21 (Fig. 5A-B and 5D), but also increased the binding of p21 to PCNA (Fig. 3C). Since several MMR components, such as MSH3, MSH6 (which recruit MSH2 to PCNA) and LIG1, interact with PCNA through their PIP motifs ^39,49^, and since the PIP box of p21 displays the highest measured affinity for PCNA ^49–51^, we speculated that the inhibitory effect of D-type cyclins on the interaction between PCNA and MMR proteins (Fig. 3A-C) is the result of p21 stabilization. In agreement with this hypothesis, overexpression of p21, but not p21(ΔPCNA), competed with MMR components for the binding to PCNA (Fig. 5E and Fig. S5I; compare lanes 3, 4, and 5), indicating a direct effect of p21 on PCNA interactors.

We propose that D-type cyclins protect p21 from degradation by shielding it from its E3 ligases. This results in increased p21 levels that, in turn, limit the binding between PCNA and the MMR components.

### Degradation of cyclin D1 in S phase is necessary to limit mutational burden and microsatellite instability

Upon S phase entry, the bulk of D-type cyclins and p21 are quickly degraded ^1,3,6,7,47,48^, while, simultaneously, MMR activity increases ^52^. Our results reveal an inhibitory role for D-type cyclins and p21 on MMR, therefore, we asked whether the cell lowers the levels of these proteins to allow adequate MMR activity during DNA replication. In contrast to parental cells, *AMBRA1*^-/-^ cells have high levels of D-type cyclins and p21 during S phase ^42^ (Fig. S6A). We observed that S phase *AMBRA1*^-/-^ cells displayed reduced binding between PCNA and MMR components in the absence of any genotoxic stress (Fig. 6A and Fig S6B; left panels, compare lane 2 to lanes 3 and 4). A reduced presence of MMR proteins was also evident in the chromatin fraction (Fig. 6A, right panel; compare lanes 1 and 2 to 3 and 4). Accordingly, compared to parental cells, *AMBRA1*^-/-^ cells displayed reduced MMR activity in S (Fig. 6B). S phase *AMBRA1*^-/-^ cells also displayed a reduced accumulation of MSH2, MSH3, and MSH6 at laser-induced DNA damage sites (Fig. 6C). Furthermore, *AMBRA1*^-/-^ cells displayed reduced binding between PCNA and MMR components upon oxidative DNA damage in S (Fig. 6D and Fig S6C; left panels, compare lane 3 to lanes 4 and 5).

**Figure 6.**
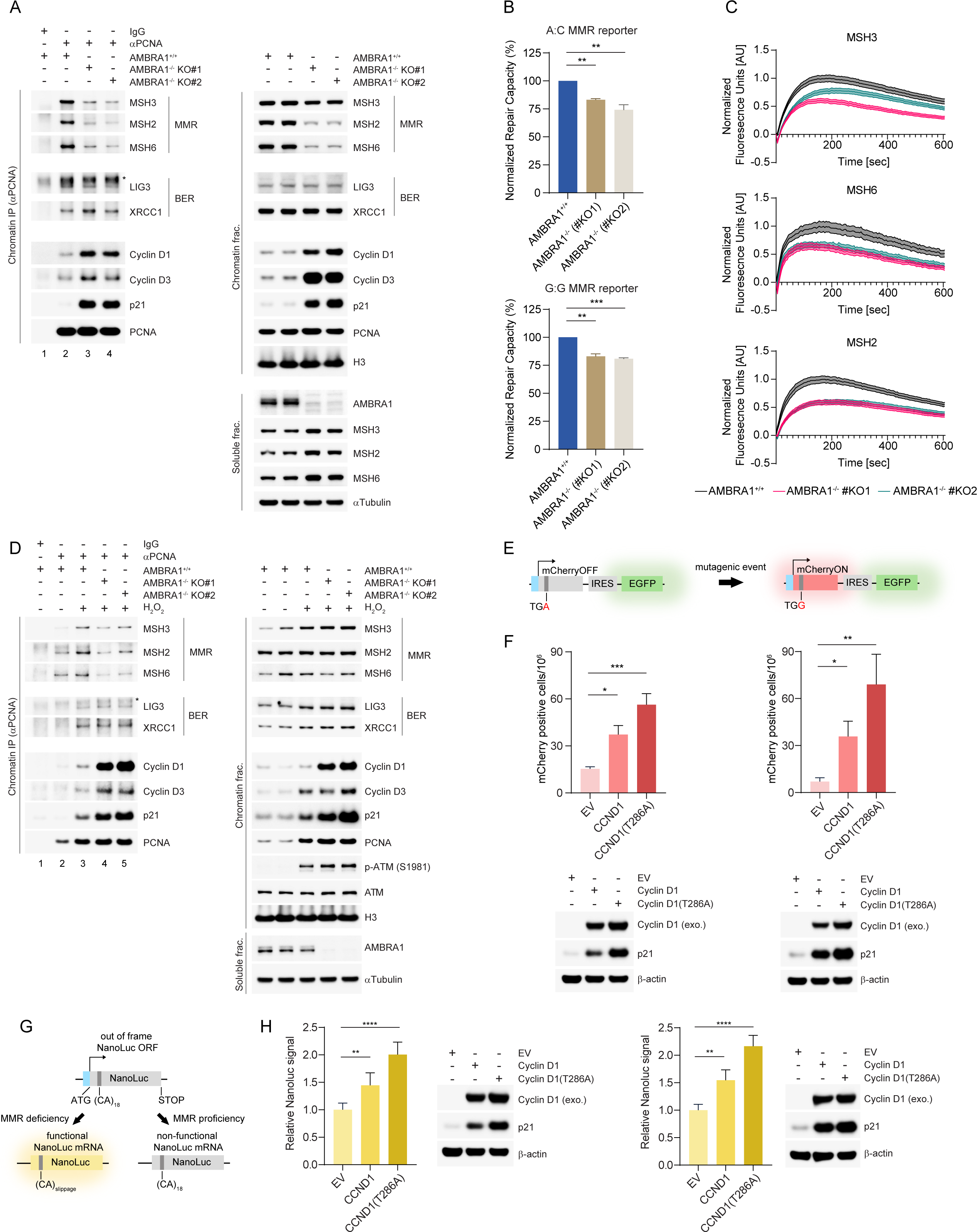
D-type cyclin degradation in S phase is necessary to limit mutational burden and microsatellite instability. A) S phase-synchronized parental (*AMBRA1*^+/+^) or *AMBRA1*^-/-^ RPE1 cells (clones #KO1 and #KO2) were fractionated into soluble and chromatin fractions. PCNA was immunoprecipitated from the chromatin fraction and co-purified proteins were immunoblotted as indicated (left panel). Right panels show chromatin and soluble fractions. Asterix denoted a non-specific band. B) S phase-synchronized parental (*AMBRA1*^+/+^) or *AMBRA1*^-/-^ RPE1 cells (clones #KO1 and #KO2) were nucleoporated with the indicated FM-HCR reporter plasmids. Eight hours later, the cells were subjected to FACS analysis. DNA repair activity for each sample was normalized to *AMBRA1*^+/+^ cells and is represented as normalized repair activity. Graphs show average and standard deviation from at least 3 independent experiments. C) DNA damage recruitment dynamics were captured by live-cell imaging in S phase-synchronized parental (*AMBRA1*^+/+^) or *AMBRA1*^-/-^ U-2OS cells (clones #KO1 and #KO2) stably expressing AcGFP-tagged MSH2, MSH3 and MSH6. Relative fluorescence values and images were acquired every 5 seconds for 10 minutes. Mean relative fluorescence values and standard errors were plotted against time. Times are indicated in seconds. D) S phase-synchronized parental (*AMBRA1*^+/+^) or *AMBRA1*^-/-^ RPE1 cells (clones #KO1 and #KO2) were treated with 100 μM H_2_O_2_ before being fractionated into soluble and chromatin fractions. PCNA was immunoprecipitated from the chromatin fraction and co-purified proteins were immunoblotted as indicated (left panel). Right panels show chromatin and soluble fractions. Asterix denoted a non-specific band. E) Schematic representation of the CherryOFF reporter assay. The CherryOFF reporter expresses two fluorescence proteins, mCherry and EGFP, driven by a CMV promoter (blue) and separated with an internal ribosome entry site (IRES). The mCherry contains a point mutation, converting its Trp98 residue (TGG) into a premature STOP codon (TGA). The resulting protein does not have fluorescence. Only a reversion of A to G in the sense strand or a T to C reversion in the antisense strand restores a functional mCherry ORF, allowing its expression. F) Quantification of the mutational frequencies in U-2OS (left) and MCF10Am (right) cells measured with the mCherryOFF reporter. mCherry signal was evaluated 18 days after infecting the cells with lentiviruses expressing either the stable cyclin D1(T286A) mutant, wild-type cyclin D1, or an empty vector (EV). Graphs show average and standard error from at least 3 independent experiments. Cyclin D1 and p21 levels are shown in the panels below the graphs. G) Schematic representation of the (CA)_18_-NanoLuc reporter assay. The CMV promoter-driven (blue) NanoLuc enzyme is out-of-frame due to a microsatellite region, composed of 18 (CA)-dinucleotide repeats. In the absence of proper MMR, random mutations that introduce frameshifts in this region can restore the NanoLuc ORF, leading to luminescence. H) Graphs show NanoLuc signal in U-2OS (left) and MCF10Am (right) cells normalized to cell viability. NanoLuc signal was evaluated 144 hours after infecting the cells with lentiviruses expressing either the stable cyclin D1(T286A) mutant, wild-type cyclin D1, or an empty vector (EV). Graphs show average and standard deviation from at least 3 independent experiments. Cyclin D1 and p21 levels are shown in the panels next to the graphs.

The experiments in Fig. 6A-D suggest that elevated levels of D-type cyclins inhibit MMR activity. Upon partial ablation of MMR proteins, changes in cellular MMR activity of a magnitude that was similar to those observed in our experiments, led to profound cellular effects, such as enhanced sensitivity to alkylating agents and increased mutational burden^45,46^. Thus, to further evaluate the functional effects of elevated cyclin D1 levels, we used two different reporter systems to measure the mutation frequency of cells overexpressing cyclin D1 as compared to parental cells. The first one, the mCherryOFF reporter, relies on a gain-of-fluorescence approach where spontaneous mutagenic events can restore the open reading frame of mCherry that contains a premature stop codon ^53^ (Fig. 6E). We expressed either the stable cyclin D1(T286A) mutant, wild-type cyclin D1, or an empty vector (EV) in U-2OS and MCF10Am cells. As expected, cyclin D1 overexpression increased p21 levels in both cell lines (Fig. 6F). Cyclin D1(T286A) and, to a lesser extent, wild-type cyclin D1 elevated random mutagenic events as measured by the increase in the fraction of red cells (Fig. 6F and Fig S6D). Next, we used an assay that is particularly sensitive to MMR activity. This assay is based on the use of the (CA)_18_-NanoLuc reporter encoding the NanoLuc enzyme that is out-of-frame due to a microsatellite region containing 18 (CA)-dinucleotide repeats. In the absence of proper MMR, random mutations that introduce frameshifts in this region can restore the NanoLuc ORF, leading to luminescence ^46^ (Fig. 6G). As expected, in both U-2OS and MCF10Am cells, silencing of both MSH2 and MLH1 lead to an increase in the NanoLuc signal compared to a non-targeting control siRNA (Fig. S6E-F). In cells expressing cyclin D1(T286A) and to a lesser extent wild-type cyclin D1, the NanoLuc signal was also significantly higher compared to EV infected cells (Fig. 6H). This difference suggests that elevated levels of cyclin D1 reduces MMR capability, leading to elevated mutation frequency.

Taken together, our results highlight the relevance of D-type cyclins degradation during S phase to maintain genomic integrity.

## DISCUSSION

The major function of the MMR pathway is to correct mismatched nucleotides and small insertion/deletion loops (IDLs) during the S phase of the cell cycle ^52,54^. MMR is also active in G1 ^15–18^, in the absence of DNA replication, to process abnormal base pairs derived from errors made by other DNA repair pathways during repair synthesis, as well as a variety of abnormal base pairs resulting from DNA damage (*e.g.*, lesions containing O^6^-methylguanine, 8-oxoguanine, carcinogen adducts, UV photo products, and cisplatin adducts) ^19,21,55^. Since oxidized bases are one of the most common DNA lesions, it is not surprising that several DNA repair pathways (*e.g.*, BER, NER, and MMR) may recognize them; however, it is not known how G1 cells decide which of these pathways to utilize for DNA repair ^19–25^. MMR induces much longer resection tracts (see Introduction) as compared to BER (*i.e.,* 1-2 bases resected in the short-patch BER and up to ∼10 bases resected in the long-patch BER) ^56–58^. This, together with lack of stand discrimination, makes MMR more likely to introduce mistakes in G1 cells and, for this reason, BER is preferred over MMR ^54^. Moreover, in G1, when there is paucity of free nucleotides, longer DNA resection can lead to fragile single-stranded DNA tracts and slower repair^15^. Consistent with the notion that MMR is kept in check during G1, we observed that inhibition of the BER pathway, but not the MMR pathway, results in a significant decrease in DNA synthesis repair in response to oxidative DNA damage (Fig. 4C-D and S4C). These results confirm that cells mainly utilize BER for repairing oxidative DNA damage during G1 ^17^. However, in G1 cells that were depleted of D-type cyclins or p21, we observed enhanced MMR activity, increased MMR-dependent DNA synthesis repair, slower recovery, and increased apoptosis (Fig. 4A-E and S4A-B). It appears that D-type cyclins exert their effects on pathway choice mainly by preventing the interaction of MMR proteins with PCNA, rather than promoting the recruitment of BER components to PCNA (Fig. 3A-D, Fig. 6A, and Fig. 6D). In other terms, during G1, BER is the default repair system of oxidative lesions, while MMR is suppressed by D-type cyclins.

Numerous studies ^59–62^ have established the molecular mechanisms dictating pathway choice between HRR and non-homologous DNA end joining. However, how the cell chooses between MMR and BER to repair DNA damage in G1 had remained unknown. Here, we describe the molecular mechanisms that inhibit MMR activity upon oxidative DNA damage in G1. Specifically, we show that D-type cyclins are recruited to sites of oxidative DNA damage in a PCNA- and p21-dependent manner (Figures 1-2 and Figures S1-S2). D-type cyclins stabilize p21, which then competes through its PIP motif with MMR components in binding to PCNA (Fig. 5). The role of D-type cyclins is crucial since p21 contains a PIP degron ^47,63^ and would otherwise be degraded if it were not protected by its binding to these cyclins. Together, these findings also provide a biological significance to the >30 years old observation that D-type cyclins and PCNA can be found in the same protein complex.

Considering PCNA’s strong involvement in almost every step of the MMR process, disrupting the interaction between MMR proteins and PCNA negatively affects the entire MMR pathway. PCNA interacts with the MutSα (MSH2/MSH6) and MutSβ (MSH2/MSH3) complexes to help recognize mismatches and IDLs, while also promoting their recruitment ^64^. PCNA also activates the endonuclease activity of the MutLα complex (MLH1/PMS2) ^56,58,65,66^, regulates EXO1’s activity for resection ^67^, and recruits POLD for DNA repair synthesis ^68,69^. Despite the presence of PIP motifs in some components of the BER pathway, PCNA has a much more limited function during both short-patch and long patch BER ^70^. Accordingly, D-type cyclins co-localize with PCNA and BER factors at oxidative DNA damage sites (Fig. 2H).

In S phase, MMR displays its highest activity ^52^ despite little fluctuations of MMR protein levels at G1/S ^71–74^. Thus, the reason for this increased activity was not fully understood. Our findings suggest that the CRL4^AMBRA1^-dependent degradation of D-type cyclins makes p21 susceptible to proteolysis via CRL1^SKP2^ and CRL4^CDT2^ (Fig. 5). In turn, this event enables proper binding of MMR proteins to PCNA (Fig. 6) to address DNA replication errors. We show that failure to degrade D-type cyclins at G1/S and the consequent p21 stabilization reduce MMR activity (Fig. 6), explaining the observation that cyclin D1 expression is barely detectable in the nucleus of S phase cells ^1,3,4,6,7^, unless AMBRA1 is knocked out ^42^. In light of our findings, it is possible that overexpression of D-type cyclins, which is often observed in cancer cells (*e.g.*, due to amplification of their loci, mutations in their degrons, as well as low levels or mutations in *AMBRA1*), negatively affects MMR, contributing to genome instability. Accordingly, cyclin D1 overexpression increases mutational burden and microsatellite instability (Fig. 6E-H). This could not be equally achieved by overexpression of p21 since p21 also functions as a tumor suppressor by inhibiting CDK2 and, consequently, cell proliferation. As the effect of D-type cyclins on MMR is CDK-independent, the present study provides insights into the challenges of therapies with CDK4/6 inhibitors both in the clinic (in breast cancer patients) and in ongoing clinical trials.

In contrast to cyclin A2 and cyclin E that, upon DNA damage in S and G2, participate to HRR by promoting DNA resection ^8^, here we suggest that during G1, D-type cyclins limit the processing of oxidative DNA lesions by MMR, reducing DNA resection without affecting BER (see model in Fig. 7). Moreover, we propose that the timely degradation of D-type cyclins is necessary for proper MMR function in S phase (Fig. 7). Intriguingly, cyclin D1 has been shown to be present in quiescent ^4,75,76^ and terminally differentiated cells ^77–81^, which represent the very large majority of cells in adult humans. However, the function of cyclin D1 expression in these non-proliferating cells remains unclear, especially given the absence of CDK activity. Thus, it is tempting to speculate that the mechanisms we have described for G1 cells also operate in both G0 and post-mitotic cells, particularly in neurons that are under the constant threat of oxidative DNA damage.

**Figure 7.**
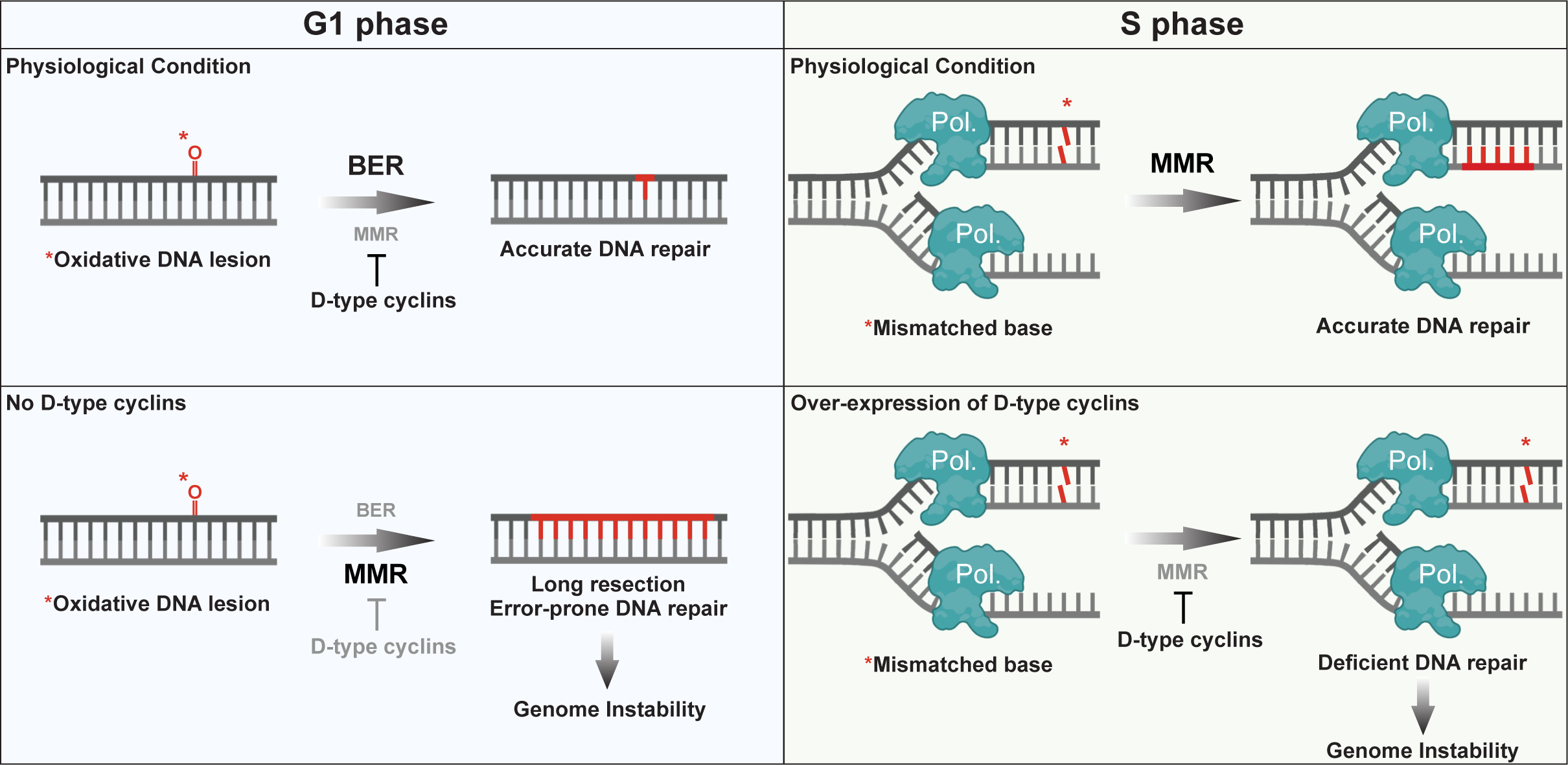
**Schematics representing how D-type cyclins regulate MMR and genome stability during the G1 and S phases of the cell division cycle.**

## Materials and Methods

### Cell culture

Cell lines were purchased from ATCC, unless otherwise stated, and were routinely checked for mycoplasma contamination with the MycoStrip™ - Mycoplasma Detection Kit (Invivogen). No cell lines used in this study were found in the database of commonly misidentified cell lines that is maintained by ICLAC and NCBI Biosample. *AMBRA1*^-/-^ U-2OS and *AMBRA1*^-/-^ RPE1 cells were previously generated ^42^. U-2OS cells harboring a stably integrated lacO array and a doxycycline-inducible transcription reporter construct at a single locus (U2OS 2-6-3 cells) were a gift from Dr. Li Lam (University of Pittsburgh, US) ^33^. hTERT RPE1 cells with mVenus-tagged CCND1, mTurquois-tagged p21 and mRuby-tagged PCNA were a gift from Dr. Jörg Mansfeld (Institute of Cancer Research, UK) ^5^. MCF10A and MCF10Am (*TP53*^-/-^ MCF10A) were purchased from Horizon Discovery. TK6 cells (wild type and MLH1-knockout), reported previously ^82^, were purchased from the TK6 Consortium in Kyoto and cultured in RPMI medium with 10% FBS. HEK 293T (ATCC CRL-3216) and hTERT RPE1 (ATCC CRL-4000) cells were maintained in Dulbeccòs modified Eaglès medium (Gibco), and U-2OS (ATCC HTB-96) cells were maintained in McCoy’s 5A medium (Gibco). Media was supplemented with 10% fetal bovine serum (FBS) (Corning Life Sciences) and 1% penicillin/streptomycin/L-glutamine (Corning Life Sciences). MCF10A and MCF10Am cells were cultured in DMEM:F12 mixture (Gibco) supplemented with 5% horse serum (Gibco), 1% penicillin/streptomycin (Corning Life Sciences), 100 ng/ml cholera toxin (Sigma-Aldrich), 10 μg/ml human insulin (Sigma-Aldrich), 10 ng/ml recombinant human EGF (Sigma-Aldrich) and 0.5 μg/ml hydrocortisone. U-2OS cells stably infected with FLAG-CCND1/2xStrep-CCND2/HA-CCND3 were maintained in McCoy’s 5A medium supplemented with 10% Tet System Approved FBS (Takara, Clontech Laboratories) and 1% penicillin/streptomycin/L-glutamine (Corning Life Sciences). All cell lines were maintained at 37°C and 5% CO2 in a humidified atmosphere.

### Plasmids, siRNA and transfection

*Homo sapiens* cDNAs were amplified by PCR using Q5 High Fidelity DNA Polymerase (New England Biolabs) and sub-cloned into a variety of vector backbones as indicated in the figure legends. This includes modified pcDNA3.1 vectors containing N-terminal FLAG/HA- or Strep tags; pBABE.puro/hygro retroviral vectors containing N-terminal FLAG/HA/Strep/mAzGreen/EGFP/mCherry/mPlum/mCerulean or TurboID-tags; pRetroQ.puro retroviral vectors containing either N- or C-terminal mCherry/AcGFP or mNeonGreen-tags; or pLVX-IRES-puro, pLVX-IRES-mCherry, pTRIPZ.puro or pLV lentiviral vectors containing N-terminal FLAG/Strep/HA/mCherry/EGFP or TurboID-tags. Specific details will be provided on request. FFSS indicates a tandem 2×FLAG-2×Strep tag; SF indicates a Strep-FLAG tag. Site-directed mutagenesis was performed using Q5 High Fidelity DNA Polymerase (New England Biolabs). The NanoLuc expressing plasmid was provided by PhoreMost Ltd (Cambridge, UK). All cell lines were transiently transfected using Lipofectamine 3000 (ThermoFisher Scientific) based on the manufacturer’s recommendation. siRNA oligo transfections were performed using RNAiMax (ThermoFisher Scientific) according to the manufacturer’s instructions.

The following siRNA duplexes from Dharmacon were used:

CCND1: L-003210-00-0005,
CCND2: L-003211-00-0005,
CCND3: L-003212-00-0005,
CDKN1A: L-003471-00-0005,
CDKN1B: L-003472-00-0005,
CDKN1C: L-003244-00-0005,
CDK4: L-003238-00-0005,
CDK6: L-003240-00-0005,
ERCC1: L-006311-00-0005,
MLH1: L-003906-00-0005,
MSH2: L-003909-00-0005,
POLB: L-005164-00-0005,
TP53BP1: L-003548-00-0005,
XPA: L-005067-00-0005,
XRCC1: L-009394-00-0005.

The following plasmids from Addgene were used:

Cherry-LacI, Plasmid #18985
3xHA-TurboID-NLS, Plasmid #107171
pQC-CherryOFF-GFP, Plasmid #129101
pQC-mCherryFP-GFP, Plasmid #129102
pSpCas9(BB)-2A-GFP (PX458), Plasmid #48138
psPAX2, Plasmid #12260
pCMV-VSV-G, Plasmid #8454
pLenti CMV rtTA3 Hygro, Plasmid #26730

### Transfections and virus-mediated gene transfer

For retrovirus production, HEK 293T cells were transfected with pBABE or pRetro-Q vectors containing the genes of interest, in combination with pCMV-Gag-Pol and pCMV-VSV-G plasmids. For lentivirus production, HEK 293T cells were transfected with pLVX, pTRIPZ or pLV vectors containing the genes of interest, in combination with pxPAX2 and pCMV-VSV-G. The virus-containing medium was collected 72 h after transfection, filtered using 0.45-μm sterile Millex-HV filter units (Millipore Sigma), and supplemented with 8 μg/ml polybrene (Sigma). Cells were infected by replacing their culture medium with the virus-containing medium for 8 h. Selection of transduced cells was carried out using either 1-2 μg/ml puromycin (Sigma), 5-10 μg/ml blasticidin S (Invivogen) or FACS, depending on the selection marker.

### CRISPR–Cas9 genome editing

CRISPR–Cas9 genome editing techniques were carried out as previously described in ^42^. In brief, to generate *CCND1*, *CCND2*, *CDKN1A* knockout cells, optimal gRNA target sequences closest to the start codon of the genes were designed using the Benchling CRISPR Genome Engineering tool (https://www.benchling.com). For transient Cas9 expression, gRNAs specific for each gene were cloned into the pSpCas9(BB)-2A-GFP (PX458) vector. The following oligos were used to generate the proper gRNA in the vector: *CCND1* (F: 5’-CACCGGTTGGCATCGGGGTACGCG-3’, R: 5’-AAACCGCGTACCCCGATGCCAACC-3’), *CCND2* (F: 5’-CACCGCGGAAGGTAGCGCTCCTCGA-3’, R: 5’-AAACTCGAGGAGCGCTACCTTCCGC-3’), *CDKN1A* (F: 5’-CACCGATGTCCGTCAGAACCCATG-3’, R: 5’-AAACCATGGGTTCTGACGGACATC-3’). Cells were seeded into 10-cm dishes at approximately 70% confluency and transfected with 5 μg of gRNA-containing PX458 plasmid, using Lipofectamine 3000 (Life Technologies). Two days after transfection, GFP-positive cells were sorted using the Beckman Coulter MoFlo XDP cell sorter (100 μm nozzle), and 15000 cells were plated on a 15-cm dish. 10 days later, single-cell clones were picked, dissociated in trypsin-EDTA (Sigma-Aldrich) for 5 min, and plated into individual wells of a 96-well plate for genotyping. Genomic DNA was collected using QuickExtract (Epicentre). Genotyping PCRs were performed with MyTaq HS Red Mix (Bioline), using primers surrounding the genomic target sites. The following primers were used for genotyping: *CCND1* (F: 5’-GAAGTTGCAAAGTCCTGGAGCC-3’, R: 5’-AGGGAAGTCTTAAGAGAGCCGC-3’), *CCND2* (F: 5’-AAAAACCTTTTTCCAGGCCGGG-3’, R: 5’-ATTGCCCTCTCCCAGGTTTAGG-3’), *CDKN1A* (F: 5’-CCTGTGGGAAGGAAGCAGGA-3’, R: 5’-CGAAGTTCCATCGCTCACGG-3’). The resulting PCR products were then purified and sequenced to determine the presence of an insertion or deletion event. Clones positive for insertion or deletion events leading to frame shift and therefore gene disruption, were validated by western blot.

### Antibodies

The following antibodies were used:

β-actin (1:5000, Sigma-Aldrich A5441)
AMBRA1 (1:1000, Cell Signaling Technology 24907S)
ATM (1:1000, Cell Signaling Technology 2873S)
CDK2 (1:1000, Bethyl Laboratories A301-812A)
CDK4 (1:1000, Proteintech 11026-1-AP)
CDK4 (1:1000, Santa Cruz Biotechnology sc-260)
CDK6 (1:1000, Proteintech 19117-1-AP)
CDK6 (1:1000, Santa Cruz Biotechnology sc-7961)
CDT1 (1:1000, Cell Signaling Technology 8064S)
CDT2 (1:1000, Bethyl Laboratories A300-948A)
CHK2 (1:1000, Cell Signaling Technology 3440S)
Cleaved Caspase-3 (1:1000, Cell Signaling Technology 9661S)
Cleaved PARP1 (1:1000, Cell Signaling Technology 5625S)
cyclin A (1:1000, Santa Cruz Biotechnology sc-271682)
cyclin A (1:5000, M.P. laboratory)
cyclin B1 (1:5000, Santa Cruz Biotechnology sc-245)
cyclin D1 (1:1000, Abcam ab16663)
cyclin D1 (1:1000, Santa Cruz Biotechnology sc-20044)
cyclin D2 (1:1000, Cell Signaling Technology 3741S)
cyclin D3 (1:1000, Proteintech 26755-1-AP)
cyclin D3 (1:1000, Thermo Fisher Scientific MA5-12717)
cyclin E (1:1000, Santa Cruz Biotechnology sc-247)
ERα (1:1000, Santa Cruz Biotechnology sc-8002)
ERCC1 (1:1000, Proteintech 14586-1-AP)
FLAG (1:2000, Sigma-Aldrich F1804)
FLAG (1:2000, Sigma-Aldrich F7425)
GAPDH (1:5000, Cell Signaling Technology 97166S)
HA (1:2000, Bethyl Laboratories A190-108A)
HA (1:5000, Covance MMS-101P)
Histone H3 (1:10000, Abcam ab1791)
LIG1 (1:1000, Proteintech 18051-1-AP)
LIG3 (1:1000, Proteintech 26583-1-AP)
MLH1 (1:1000, Cell Signaling Technology 3515S)
MLH1 (1:1000, Cell Signaling Technology 4256S)
Mouse Gamma Globulin Control (ThermoFisher Scientific 31878)
MSH2 (1:1000, Cell Signaling Technology 2017S)
MSH2 (1:1000, Cell Signaling Technology 2850S)
MSH3 (1:1000, Proteintech 22393-1-AP)
MSH6 (1:1000, Cell Signaling Technology 12988S)
MSH6 (1:1000, Cell Signaling Technology 5424S)
OGG1 (1:1000, Novus Biologicals NB100-106)
p21 (1:1000, Cell Signaling Technology 2947S)
p21 (1:1000, Proteintech 28248-1-AP)
p27 (1:1000, BD Biosciences 610241)
p27 (1:1000, Cell Signaling Technology 3686S)
p57 (1:1000, Cell Signaling Technology 2557S)
p57 (1:1000, Proteintech 66794-1-Ig)
PARP1 (1:1000, Cell Signaling Technology 9542S)
pATM (S1981) (1:1000, Cell Signaling Technology 4526L)
pCHK2 (T68) (1:1000, Cell Signaling Technology 2661S)
PCNA (1:2000, Cell Signaling Technology 13110S)
PCNA (1:2000, or 10 μg for IP, Thermo Fisher Scientific 14-9910-37)
pH3 (S10) (1:2000, Cell Signaling Technology 3377S)
POLB (1:2000, Proteintech 18003-1-AP)
pRB (S807/811) (1:1000, Cell Signaling Technology 8516S)
RB (1:1000, Cell Signaling Technology 9309S)
RB (1:1000, Cell Signaling Technology 9313S)
SKP2 (1:1000, Cell Signaling Technology 2652S)
TP53BP1 (1:2000, Abcam ab36823)
XPA (1:1000, Abcam ab65963)
XRCC1 (1:1000, Cell Signaling Technology 76998S)
XRCC1 (1:1000, Proteintech 21468-1-AP)
αTubulin (1:5000, Sigma-Aldrich T6074)
γH2A.X (1:1000, Cell Signaling Technology 9718S)
γH2A.X (1:1000, EMD Millipore 05-636)

### Drug treatment procedures

Where indicated, cells were treated with 0.2 ug/ml neocarzinostatin (NCS) (Sigma-Aldrich, N9162-100UG) for 30 minutes, 20 μg/ml bleomycin (Selleck, S1214) for 3 hours, 40 J/m^2^ UVC (254 nm) for 15 minutes, 50-250 μM H_2_O_2_ (Sigma-Aldrich, H1009-100ML) for 20 minutes, 500 μM MMS (Sigma-Aldrich, 129925-5G) for 1 hour, 10 μM methylnitronitrosoguanidine (MNNG) (TCI, M0527) and 10 μM O^6^-benzylguanine (Sigma-Aldrich, B2292-50MG) for 2 hours, 1 μM CPT (EMD Millipore, 208925-50MG) for 1 hour, 100 μg/ml cycloheximide (Sigma-Aldrich, C7698-1G), 1 μg/ml doxycycline (Sigma-Aldrich, D9891-1G), 10 μM EdU (Thermo Fisher Scientific, 1149-100), 10 μM MG132 (Peptides International, IZL-3175v) and 2.5 μM MLN4924 (Active Biochem, A-1139), 1 μM Palbociclib (Selleck Chemicals).

### Cell synchronization

U-2OS cells were synchronized using nocodazole. Cells were treated with 40 ng/ml nocodazole for 14 hours. Mitotic cells were then collected by shaking them off the dish, washed 3× with PBS, and replated in normal medium to allow them to resume cell cycle. Representative cell cycle progression is shown in Fig. S6A after synchronization. G1 phase experiments were carried out between 4-12 hours after nocodazole release, while S phase experiments were carried out between 20-24 hours after nocodazole release. Cells were transfected as indicated with siRNAs 8 hours before the addition of nocodazole. RPE1 cells were synchronized by serum starvation. Subconfluent plates of cells were trypsinized and washed 3× with PBS, before they were replated in DMEM containing 0.05% FBS. Cells were kept in this medium for 72 hours before they were trypsinized and replated at ∼70% confluency in DMEM containing 10% FBS to allow them to enter the cell cycle. Representative cell cycle progression is shown in Fig. S6A after serum release. G1 phase experiments were carried out between 6-16 hours after serum release, while S phase experiments were carried out between 18-24 hours after serum release. Cells were transfected as indicated with siRNAs during the 72 hours of serum starvation period.

### qRT-PCR

Total RNA was purified using RNeasy mini kits (Qiagen). cDNA was generated using Double Primed EcoDry kits (Takara). The qPCR reaction was carried out using PowerUp SYBR Green (Applied Biosystems) with the Applied Biosystems QuantStudio 3 Real-Time PCR system in a 96-well format. ROX was used as a reference dye for fluorescent signal normalization and for well-to-well optical variations correction. Bar graphs represent the relative ratios of target genes to GAPDH housekeeping gene values. For each biological sample, triplicate reactions were analyzed using absolute relative quantification method alongside in-experiment standard curves for each primer set to control for primer efficiency. The oligos used for qRT-PCR analysis were: *GAPDH* (F: 5’-TGCACCACCAACTGCTTAGC-3’, R: 5’-GGCATGGACTGTGGTCATGAG-3’), *CDKN1A* (F: 5’-CGGCAGACCAGCATGACAGATTT-3’, R: 5’-GTGGGCGGATTAGGGCTTCC-3’).

### Laser induced DNA damage and Live-Cell Imaging

Laser-induced DNA damage induction and live-cell imaging were essentially performed as in ^83^. Briefly, ∼50,000 cells were plated per well on a four-well Lab-Tek II chambered number 1.5 borosilicate coverglass for 24 hours before imaging. RNAi transfections were performed approximately 2 days prior to microscopy using two rounds of silencing. Cells were pre-sensitized with 10 μM BrdU (Merck) for 24 hours. For imaging, on the day of data collection, cells were incubated in FluoroBrite^TM^ DMEM supplemented with 10% FBS, 25 mM HEPES (Sigma-Aldrich), and 1% sodium pyruvate (Gibco). Imaging was performed using a DeltaVision Elite inverted microscope system (Applied Precision), using a 60× oil objective (1.42 NA) from Olympus. Excitation was achieved with a 7 Color Combined Insight solid state illumination system equipped with a polychroic beam splitter with filter sets to support GFP (ex.: 475/28 nm, em.: 525/50 nm) and mCherry (ex.: 575/25 nm, em.: 632/60 nm). Images were acquired using a CoolSNAP HQ2 camera. DNA damage was generated using a 405-nm, 50-mW laser at 100% power for 0.5 s. One pre-laser image was recorded, and the number and interval of post-laser images varied in each experiment (as indicated on the kinetic plots). Recruitment intensity was analyzed using a macro written for ImageJ that calculated the ratio of intensity of a defined laser spot *A* to the adjacent area *B* after subtraction of the background intensity of an unpopulated area of the image *C* ^83^. Thus, the relative fluorescent unit (RFU) for each data collection point was calculated by the equation *RFU* = (*A − C*)/(*B − C*). In instances where fluorescent recruitment was not detected, the coordinates of *A* were determined by use of laser coordinates recorded in the data log file. The mean values and standard errors from several cells and live-cell imaging time courses were computed for each time point using GraphPad Prism software.

To visualize the recruitment of endogenous cyclin D1, D2 and D3, laser induced DNA damage stripes were done on a Zeiss LSM 800 microscope, using a 405 nm diode laser (5 mW) with the timed bleach option (60 iterations, 80% laser power output) in the ZenBlue 2.1 software using a Plan-Apochromat 63×/1.40 oil objective. Cells were pre-sensitized with 1 μg/ml Hoechst 33342 for 30 minutes.

When micro-irradiation was used under specific conditions that do not induce DSBs, but generate oxidized bases, the experiments were essentially performed as in ^34^. This was used to visualize the recruitment dynamics to oxidative DNA damage sites of endogenous, mVenus-tagged cyclin D1 and mRuby-tagged PCNA or mCherry/AcGFP/mNeonGreen-tagged BER and MMR proteins. Imaging was performed using a Nikon A1R-HD25 confocal microscope, using a 60x oil plan-apochromat objective (1.42 NA). DNA damage was generated using a Nikon LUN-F 50 mW 405 nm FRAP laser unit at 100% power output with a 1000 µs dwell time. Exact optical configuration and laser setting are described in ^34^.

### Image analysis of BER and MMR protein recruitment

The average mean intensities of the micro-irradiated region and background nuclear region were calculated at five-second intervals on a 10-minute timescale using a custom multi-step ImageJ macro. In short, large fields of view containing numerous micro-irradiated cells were cropped for individual cells, and the cropped videos were registered to correct for cell movement. The nuclear region-of-interest was determined by applying an automatic threshold to each frame and the mean intensity of the background (nuclear region with stripe region removed) and stripe region over the length of the movie was measured for each cell and saved to a text file for further processing. Detailed usage instructions and macro code can be accessed at https://github.com/FenyoLab/recruitment_panel. Next, the mean intensity of the background region was subtracted from that of the micro-irradiated region, and this difference was normalized for differences in nuclear intensity by subtracting the original difference between both regions prior to micro-irradiation from all subsequent timepoints. Statistical analysis: The mean intensity of the normalized difference over sliding windows of 60 seconds across the 10-minute timescale was calculated for each nucleus in both control and silencing/knockout conditions. Then, Mann Whitney U tests for statistical significance were performed for each sliding window.

Heatmap generation: the value at each time point was divided by the maximum recruitment value across all silencing or knockout conditions for each DNA damage repair protein, resulting in standardized values for all proteins ranging from 0 to 1. Then, the mean value at each time point over all cells measured for control condition was subtracted from the mean for silencing condition. These values were averaged over 15 second intervals and a heatmap generated with the python package seaborn. Hierarchical clustering of rows in the heatmap was performed with Euclidean distance measure and Ward clustering method.

### Immunoprecipitation (IP), immunoblotting (IB) and fractionation

When no cellular fractionation was carried out, cells were lysed in lysis buffer (25 mM Tris.HCl pH 8.0, 150 mM NaCl, 0.2% NP-40, 10% glycerol, 1 mM EDTA, 1 mM EGTA, 2 mM MgCl_2_, 1 mM 1,4-Dithiothreitol (DTT) and 5 mM N-Ethylmaleimide (Sigma-Aldrich)). The insoluble fraction was removed by centrifugation (20000 x g x 15 min at 4 °C). Where indicated, cells were fractioned into soluble, and chromatin bound fractions by first lysing them in CSK buffer (10 mM PIPES pH=7.0, 100 mM NaCl, 300 mM sucrose, 0.1 % Triton X-100, 3 mM MgCl_2_, 1 mM EGTA, 1 mM DTT and 5 mM N-Ethylmaleimide) for 5 minutes and then pelleting them (3000 x g x 3 min at 4 °C). The collected supernatant corresponds to the soluble fraction. Cell pellets were then washed in CSK buffer and were subsequentially lysed in chromatin lysis buffer (50 mM TRIS.HCl pH=7.5, 150 mM NaCl, 0.2 % NP-40, 10% Glycerol, 1 mM EDTA, 1 mM EGTA, 4 mM MgCl_2_, 0.5 mM DTT, 5 mM N-Ethylmaleimide and 5 U/mL TurboNuclease (Accelagen)) for 30 minutes. The insoluble fraction was removed by centrifugation (20000 x g x 15 min at 4 °C). All of the above indicated buffers were supplemented with protease (Complete™ ULTRA, Roche) and phosphatase inhibitors (PhosSTOP™, Roche). Immunoprecipitations and affinity precipitations were carried out using either FLAG-M2 magnetic beads (Sigma-Aldrich), anti-HA magnetic beads (Pierce), MagStrep “type3” XT beads (IBA) at 4°C for 2 hours. Endogenous immunoprecipitations of PCNA, were carried out for 2 h at 4 °C using anti-PCNA antibody (PC10 clone, eBioscience) mixed with Protein A/G Magnetic Beads (Pierce) at 4°C for 2 hours. Mouse IgG isotype (Thermo Fischer Scientific) was used as a negative control. Beads were extensively washed in lysis buffer and elution was carried out with 3×FLAG peptide (Sigma-Aldrich), HA peptide (Roche), Strep-Tactin XT Elution Buffer (IBA) or 1× Laemmli buffer. Immunoblotting was performed as previously described ^83^. Briefly, samples were resolved under denaturing and reducing conditions using 4%–12% Bis-Tris gels (NuPAGE) and transferred to a PVDF membrane (Immobilon-P, Millipore). Membranes were blocked with 5% nonfat dried milk, incubated with primary antibodies overnight at 4°C. After washing the membranes, secondary antibodies coupled with horseradish peroxidase were applied (Amersham-GE). Immunoreactive bands were visualized by enhanced chemiluminescence reagent (Pierce) and signal was acquired using ImageQuant LAS 4000 (GE).

### Immunofluorescence (IF) microscopy

Cells cultured on coverslips were either pre-extracted with ice-cold CSK buffer containing 0.5% Triton X-100 for 5 minutes, or directly fixed in 4% paraformaldehyde/PBS after a PBS wash. Cells were permeabilized with PBS/0.1% Triton X-100 for 15 minutes and were then blocked with blocking buffer (PBS/0.05% Triton X-100 containing 5% FBS and 3% BSA) before incubation with indicated primary antibodies diluted in blocking buffer. Alexa-488, Alexa-555 and Alexa-647-conjugated secondary antibodies were from Thermo Fischer Scientific and used at 1:2000 dilution. To identify cells undergoing DNA replication, cells were pulsed with EdU (10 μM final concentration, Thermo Fisher Scientific) for 45 minutes, washed with PBS and fixed in 4% PFA. Detection of EdU was accomplished using the Click-iT Plus EdU Alexa Fluor 488 or 647 Imaging Kit (Thermo Fisher Scientific) according to manufacturer’s instructions. Slides were mounted in ProlongDiamond with DAPI (Molecular probes). Imaging was performed using a DeltaVision Elite inverted microscope system (Applied Precision), using a 60x/1.42 NA oil objective from Olympus, a Zeiss Axio Observer using a Plan-Apochromat 63×/1.40 NA oil objective, a Zeiss LSM 800 microscope using a Plan-Apochromat 63x/1.40 NA oil objective or a Nikon A1R-HD25 confocal microscope system, using a 60x 1.4 NA oil plan-apochromat objective.

### Site-specific ROS and DSB induction

KillerRed cDNA (obtained from Dr. Li Lam, University of Pittsburgh, US), mCherry-FokI and the catalytically dead mCherry-FokI (D540A) cDNAs were subcloned into the mCherry-LacI vector (Addgene plasmid #18985) replacing mCherry. 2-6-3 U-2OS cells harboring a stably integrated lacO array and a doxycycline-inducible transcription reporter construct at a single locus were transfected with the indicated constructs. Immediately after transfection, dishes containing cells were wrapped in aluminum foil to prevent KillerRed activation. The induction of ROS by bulk activation of KillerRed was done essentially as described previously ^33^. Briefly, 24 h after transfection, cells were exposed to a white LED light source for 15 minutes (5W Slimlight transilluminator) by placing them directly on top of the transilluminator. After exposure, the cells were immediately pre-extracted with ice-cold CSK buffer for 5 minutes before fixing them with 4% PFA in PBS. Cells were then processed as described in the Immunofluorescence (IF) microscopy section. Around 40 nuclei were quantified for each independent experiment.

### Proximity labeling (TurboID-cyclin D1 and TurboID-PCNA)

For proximity labeling of cyclin D1 partners, asynchronous U-2OS cells stably expressing 3xHA-TurboID, 3xHA-TurboID-CCND1 (1×: with a ∼5 nm linker) or 3xHA-TurboID-13x-CCND1 (13x: with a ∼25 nm linker) from a pBABE.puro backbone were used ^35,36^. Two independent experiments were performed using the ∼5 nm linker and 2 independent experiments were performed using the ∼25 nm linker. Cells were incubated with 50 μM biotin in the presence or absence of 250 μM H_2_O_2_ for 1.5 hours. Cells were first lysed in CSK buffer (10 mM PIPES pH=7.0, 0.1 % Triton X-100, 100 mM NaCl, 300 mM sucrose, 3 mM MgCl_2_, 1 mM EGTA, 1 mM DTT, 5 mM N-Ethylmaleimide) for 5 minutes and then pelleted (3000 x g x 3 min at 4 °C). Pellets were washed in CSK buffer and were subsequentially lysed in a modified RIPA buffer (25 mM Tris.HCl pH 7.6, 250 mM NaCl, 1% NP-40, 1% sodium deoxycholate, 0.1% SDS, 1 mM DTT, 5 mM N-Ethylmaleimide and 5 U/mL TurboNuclease). All the indicated buffers were supplemented with protease (Complete™ ULTRA, Roche) and phosphatase inhibitors (PhosSTOP™, Roche). The insoluble fraction was removed by centrifugation (20000 x g x 15 min at 4 °C). Affinity purifications were carried out using MyOne Streptavidin C1 beads (Thermo Fisher Scientific) at 4°C for 4 hours. Beads were extensively washed in RIPA buffer and in high salt RIPA buffer (containing 0.5 M NaCl) before elution. Elution was carried out by keeping the beads at 95°C for 5 minutes in elution buffer (70 mM Tris.HCl pH=7.5, 2% SDS, 0.5 mM EDTA, 350 mM 2-mercaptoethanol). For proximity labeling of PCNA partners, G1 synchronized RPE1 cells stably expressing NLS-3xHA-TurboID or NLS-3xHA-TurboID-13x-PCNA (13x: with a ∼25 nm linker) from a pLVX.puro backbone were used. RPE1 cells transfected with either non-targeting (siNT) or siRNA targeting D-type cyclins (siCCNDs) were processed identically to the proximity labeling of cyclin D1 partners described above.

### Sample preparation prior to mass spectrometry for TurboID-cyclin D1

Affinity purified proteins were reduced with DTT at 57 °C for 1 h (2 µl of 0.2 M) and subsequently alkylated with iodoacetamide at RT in the dark for 45 minutes (2 µl of 0.5 M). To remove detergents from the chromatin fractions and to also increase the dynamic range of the analysis the samples were loaded onto a 1.0 mm NuPAGE 4-12% Bis-Tris Gel (Life Technologies). The gel was run for approximately 30 minutes and stained using GelCode Blue Stain Reagent (Thermo Scientific). The dominant histone H1 bands were excised, extracted and analyzed on the mass spectrometer separately from the remainder of the lane. The excised gel pieces were destained in 1:1 v/v solution of methanol and 100 mM ammonium bicarbonate solution using at least three exchanges of destaining solution. The destained gel pieces were partially dehydrated with an acetonitrile rinse and further dried in a SpeedVac concentrator for 20 minutes. 200 ng of sequencing grade modified trypsin (Promega) was added to each sample. After the trypsin was absorbed, 250 µl of 100 mM ammonium bicarbonate was added to cover the gel pieces. Digestion proceeded overnight on a shaker at RT. A slurry of R2 20 µm Poros beads (Life Technologies Corporation) in 5% formic acid and 0.2% trifluoroacetic acid (TFA) was added to each sample at a volume equal to that of the ammonium bicarbonate added for digestion as described previously ^84^. The digested peptides were allowed to bind to the Poros beads for 3 h with vigorous shaking at 4 °C. The peptide loaded beads were transferred onto equilibrated C18 ZipTips (Millipore) using a microcentrifuge for 30 seconds at 6000 rpm. Gel pieces were rinsed three times with 0.1% TFA and each rinse was added to its corresponding ZipTip followed by microcentrifugation. The beads were further washed with 0.5% acetic acid. Peptides were eluted by the addition of 40% acetonitrile in 0.5% acetic acid followed by the addition of 80% acetonitrile in 0.5% acetic acid. The organic solvent was removed using a SpeedVac concentrator and the sample reconstituted in 0.5% acetic acid.

### Sample preparation prior to mass spectrometry for TurboID-PCNA

Affinity purified proteins were reduced with DTT at 57 °C for 1 h (2 µl of 0.2 M) and subsequently alkylated with iodoacetamide at RT in the dark for 45 minutes (2 µl of 0.5 M) and the alkylation was stopped with addition of another aliquot of DTT. The sample was loaded onto S-TRAP™ (Protifi) and proceeded following the manufactures protocol. In brief, the samples were acidified with 12% phosphoric acid, resuspended in S-TRAP binding buffer and loaded onto the S-TRAP. The sample was washed 3 times using the S-TRAP binding buffer. 1 μg of trypsin in 50 mM Triethylammonium bicarbonate (TEAB) was added and the digestion was allowed to proceed at 47°C for one hour. Peptides were eluted by the addition of 40% acetonitrile in 0.5% acetic acid followed by the addition of 80% acetonitrile in 0.5% acetic acid. The organic solvent was removed using a SpeedVac concentrator and the sample reconstituted in 0.5% acetic acid.

### Mass spectrometry analysis for TurboID-cyclin D1

LC separation was performed online on an EASY-nLC 1000 (Thermo Scientific) utilizing Acclaim PepMap 100 (75 μm x 2 cm) precolumn and PepMap RSLC C18 (2 um, 100A x 50 cm) analytical column. Peptides were gradient eluted directly to an Orbitrap Q Exactive HF-X mass spectrometer (Thermo Fisher) using a 95 min acetonitrile gradient from 5 to 35 % B in 60 min followed by a ramp to 45% B in 10 min and 100% B in another 10 min with a hold at 100% B for 10 min (A=2% acetonitrile in 0.5% acetic acid; B=80% acetonitrile in 0.5% acetic acid). Flowrate was set to 200 nl/min. High resolution full MS spectra were acquired with a resolution of 45,000, an AGC target of 3e6, with a maximum ion injection time of 45 ms, and scan range of 400 to 1500 m/z. Following each full MS scan 20 data-dependent HCD MS/MS scans were acquired at the resolution of 15,000, AGC target of 1e5, maximum ion time of 120 ms, one microscan, 2 m/z isolation window, normalized collision energy (NCE) of 27, fixed first mass 150 m/z and dynamic exclusion for 30 seconds. Singly charged ions and ions carrying 8 or more charges were excluded from triggering an MS2 scan. Both MS and MS2 spectra were recorded in profile mode.

### Mass spectrometry analysis for TurboID-PCNA

LC separation was performed as described above, but peptides were gradient eluted from the column directly to an Orbitrap Q Exactive mass spectrometer (Thermo Fisher) using a 155 min acetonitrile gradient from 5 to 35 % B in 120 min followed by a ramp to 45% B in 10 min and 100% B in another 10 min with a hold at 100% B for 10 min (A=2% acetonitrile in 0.5% acetic acid; B=80% acetonitrile in 0.5% acetic acid). Flowrate was set to 200 nl/min. High resolution full MS spectra were acquired with a resolution of 70,000, an AGC target of 1e6, with a maximum ion injection time of 120 ms, and scan range of 400 to 1500 m/z. Following each full MS scan 20 data-dependent HCD MS/MS scans were acquired at the resolution of 17,500, AGC target of 5e4, maximum ion time of 120 ms, one microscan, 2 m/z isolation window, normalized collision energy (NCE) of 27, fixed first mass 150 m/z and dynamic exclusion for 30 seconds. Singly charged ions and ions carrying 8 or more charges were excluded from triggering an MS2 scan Both MS and MS2 spectra were recorded in profile mode.

### Mass spectrometry data analysis

The MS/MS spectra were searched against a Uniprot (www.uniprot.org) human protein database with common lab contaminants and the sequence of the tagged bait proteins added using SEQUEST within Proteome Discoverer 1.4 (Thermo Fisher). The search parameters were as follows: mass accuracy better than 10 ppm for MS1 and 0.02 Da for MS2, two missed cleavages, fixed modification carbamidomethyl on cysteine, variable modification of oxidation on methionine and deamidation on asparagine and glutamine. The data was filtered using a 1% FDR cut off for peptides and proteins against a decoy database and only proteins with at least 2 unique peptides were reported. In all data analyses, for samples where a given protein had no observed peptide spectrum matches (PSMs), the zero entry was replaced with a count of 1 (the minimum observable PSM count being 2 due to the requirement of at least 2 unique peptides per protein). All enrichment analysis were done by Fisher’s Exact Test using protein lists generated by literature curation of KEGG pathway lists (in the case of the pathways: REP, BER, NER, MMR, HRR, NHEJ, FAN, DR, TLS, and DDA) as well as a download from BioGRID circa 7/12/22 (in the case of PCNA related proteins). When p-values were reported along with fold changes (*e.g.,* in the case of Fig. S3C), they originated from Welch’s t-test uncorrected for multiple hypothesis testing. In the case of Fig. 2H the cyclin D1 fold change was based on the most extreme ratio (*i.e.,* the one with the largest absolute log fold change) observed using either of the two linker lengths (∼5 nm or ∼25 nm linker).

### Comet assay

Comet assay was performed with the CometAssay Single Cell Gel Electrophoresis Assay Kit (Trevigen), following the manufacture’s recommendations. Under the experimental conditions described here, the assay can pick up both single- and double-stranded breaks. Briefly, G1 synchronized U-2OS cells were treated where indicated with 50 μM H_2_O_2_ for 15 minutes in Live Cell Imaging Solution (Life Technologies). Cells were allowed to recover up to 4 hours in culture media after removing H_2_O_2_ from them. When indicated, cells were pre-treated with 1 μM palbociclib before treatment with H_2_O_2_ which was maintained on the cells during H_2_O_2_ treatment and the recovery phase. At the indicated time points the cells were gently trypsinized and kept in the dark for further processing. Cells were pelleted and washed with PBS before resuspending them in ice-cold PBS at 3×10^5^ cell/ml concentration. To 60 μl of LMAagarose (Trevigen), kept at 37℃, 6 ul of the cell suspension was added, and 50 μl of this mixture was immediately pipetted onto prewarmed (37℃) CometSlide (Trevigen), that is specially treated to promote adherence of low melting point agarose. Comet slides were then kept at 4℃ for 15 minutes for a clear ring to indicate the gelling of agarose on the slide. Slides were then incubated overnight in prechilled comet lysis buffer (containing 10% DMSO) (Trevigen) at 4℃. Slides were then dipped into an alkaline solution (300 mM NaOH, 1 mM EDTA in distilled water) for 15 minutes at room temperature to aid the visualization of single stranded breaks. CometAssay ES unit (Trevigen) was used for electrophoresis, filled with 950 ml of neutral electrophoresis buffer (100 mM Tris pH=9.0, 300 mM sodium acetate) for 1h at 4℃ using 21 V. Excess running buffer was drained from the slides before immersing them into the DNA precipitation solution (1 M ammonium acetate solution in ethanol) and for 30 minutes at room temperature. Excess DNA precipitation solution was drained, and slides were kept in 70% ethanol for 30 minutes at room temperature. Excess 70% ethanol was drained, and the slides were dried by storing them at 45℃ for 15 minutes. Drying brings all the cells to a single plane. 100 µl of 1× SYBR gold (diluted in 10 mM Tris-HCl pH 7.5, 1 mM EDTA) was used for staining DNA by incubating it on the slides for 30 minutes in the dark, at room temperature. Excess SYBR gold staining solution was removed, and slides were washed several times in distilled water by using a coplin jar. After air drying, the samples were imaged on a EVOS microscope (Invitrogen) using a GFP filter set and a 10x objective. Tail moment was measured using the OpenComet plugin for ImageJ ^85^. At least 40 comet tails were analyzed for each independent experiment.

### DNA repair synthesis assay (AKA unscheduled DNA synthesis assay)

G1 synchronized RPE cells were grown on poly-D-lysine (neuVitro) or Vitronectin XF (Stemcell Technologies) coated coverslips and treated, where indicated, with 200 μM H_2_O_2_ for 30 minutes in DMEM supplemented with 1% of dialyzed FBS, 10 μM EdU and 100 nM raltitrexed. Cells were allowed to recover for 1.5 hours in DMEM supplemented with 1% of dialyzed FBS, 10 μM EdU and 100 nM raltitrexed after removing the H_2_O_2_ from them. When indicated, cells were pre-treated with 1 μM palbociclib before the treatment with H_2_O_2_, which was maintained on the cells during H_2_O_2_ treatment and the recovery phase. EdU labeling was eventually quenched by changing the media to DMEM with 1% of dialyzed FBS, and 10 μM thymidine for 15 minutes. After a PBS wash, the cells were fixed with 3.6% PFA in PBS containing 0.5% Triton X-100 for 15 min at room temperature. After washing twice with PBS, cells were further permeabilized with 0.5 % Triton X-100 in PBS for 20 minutes. Cells were blocked with 5% BSA in PBS for 10 minutes before proceeding with the click-it reaction. Click-it reaction was performed using the Click-iT Plus Picolyl Azide Toolkit (Thermo Fisher Scientific) according to manufacturer’s instructions with modifications. Picolyl-Azide-PEG4-Biotin (Jena Bioscience) was used in the reaction at a final concentration of 20 μM to react with the incorporated EdU in the presence of a CuSO_4_:copper protectant mixture of 9:1, for 30 minutes. Coverslips were washed once with PBS and once with PBS containing 5% BSA before incubating it with Alexa-488 conjugated streptavidin (Thermo Fisher Scientific) at 1:2000 dilution for 30 minutes at room temperature. After several rounds of PBS wash, the coverslips were mounted with ProlongDiamond with DAPI (Molecular probes). Imaging was performed using a DeltaVision Elite inverted microscope system (Applied Precision), using a 60x/1.42 NA oil objective from Olympus, or a Zeiss Axio Observer using a Plan-Apochromat 63x/1.40 NA oil objective. Cells with above 10-fold normalized fluorescence value were excluded from the analysis as those are S phase contaminants. At least 150 nuclei were analyzed for each independent experiment.

### Fluorescence-based multiplex host cell reactivation assay (FM-HCR)

FM-HCR assay was essentially carried out as previously described in ^43,44,86^ using synchronized RPE1 cells. To measure DNA repair capacity in G1, serum-starved RPE1 cells where trypsinized and 0.5×10^6^ of the cells were resuspended in 20 μl of the P3 Primary Cell 4D-Nucleofector Kit buffer (Lonza). The cell suspension was then nucleoporated with reporter plasmids with DNA lesions and undamaged control plasmids, using the 4D-Nucleofector unit (Lonza) with the EA-104 program. Cells were then replated into DMEM containing 10% FBS. Ten hours post nucleofection, cells were harvested and analyzed by flow cytometry. Where indicated, cells were pre-treated with 1 μM palbociclib before nucleoporation, which was maintained on the cells until cells were prepared for flow cytometry. To measure DNA repair capacity in S phase, RPE1 cells were serum starved for 72 hours and released into serum containing media for 20 hours. Cells were then collected and nucleoporated as described above. 8 hours post nucleofection, cells were harvested and analyzed by flow cytometry. TK6 cells were electroporated using the MXCell transfection system, with a voltage setting of 260 V and a capacitance setting of 960 mF. Fluorescent protein signal was quantitated for each respective plasmid reporter from at least 20000 cellular events per replicate (n ≥ 3). Normalized repair capacity was calculated as previously described ^43,44^. Welch’s t-test was used for statistical analysis. A representative FACS gating strategy is provided in Fig. S4I.

### Mutagenesis frequency measurement using the mCherryOFF reporter

To measure global mutation frequency, we used the gain-of-function mutation-activated fluorescence reporter, termed mCherryOFF. The assay was performed as previously described ^53^. In brief, stable, mCherryOFF reporter containing U-2OS and MCF10Am cell lines were generated with retroviral transduction followed by FACS selection for the EGFP signal and negative selection for any spontaneous red revertants (containing a functional mCherry). The newly generated cell lines were then infected with lentiviruses carrying either an empty vector (EV), wild-type CCND1 or CCND1(T286A) for overexpression. Transduced cells were selected for two days with puromycin. Cells were then measured for green and red fluorescence using FACS after 18 days following puromycin selection. A representative FACS gating strategy is provided in Fig. S6D.

### Mutagenesis frequency measurement using the (CA)_18_-NanoLuc reporter

To measure mutagenesis rate due to lack of MMR, we used the (CA)_18_-NanoLuc reporter. The reporter plasmid was provided by PhoreMost Ltd (Cambridge, UK). The assay was performed as previously described ^46^. Briefly, stable U-2OS and MCF10Am cell lines were generated by lentiviral transduction followed by 5-7 days of blasticidin S selection. Cells were then infected with lentiviruses to express either the stable cyclin D1(T286A) mutant, wild-type cyclin D1, or an empty vector (EV). Transduced cells were selected for two days with puromycin. Four days after puromycin selection, NanoLuc expression was measured using NanoGlo® Luciferase Assay System (Promega), according to manufacturer’s instructions. Luminescence was measured by a BioTek Synergy Neo2 plate reader. The NanoLuc signal was normalized to the viability measured through AlamarBlue assay (BioRad). Where indicated (CA)_18_-NanoLuc reporter containing stable cell lines were transfected with siRNA targeting MSH2/MLH1 or a non-targeting control siRNA. After 4 days and 2 rounds of siRNA transfection, NanoLuc expression was measured as described above.

### Statistical Analysis

Data were analyzed using GraphPad Prism 9.5.1 software. Two-group datasets were analyzed using the nonparametric Mann-Whitney test unless otherwise stated. For comparisons between three or more groups, parametric or non-parametric (Kruskal-Wallis test) one-way analysis of variance (ANOVA) was used.

## Acknowledgement

The NanoLuc expressing plasmid was a kind gift from PhoreMost Ltd (Cambridge, UK) under a Material Transfer Agreement. We thank Dr. Jörg Mansfeld (Institute of Cancer Research, UK), Dr. Li Lam (University of Pittsburgh, US) and Dr. Richard Possemato (NYU School of Medicine, US) for reagents. We thank Alberto Ciccia, Anna Pluciennik, Beata G. Vertessy, Daniel Durocher, Eli Rothenberg, Jiri Lukas for critically reading the manuscript. MP is thankful to TM Thor and TB Balduur for their continuous support.

## Author contributions

GR and MP conceived the study. GR designed, carried out the experiments, and analyzed the data. For some experiments, GR was helped by BM, HVG, JBH, QZ, DS, JWH, ES, AAW, MAK, EL, SK, LSP, JA, MA, and BU. MP supervised the experiments. LX, AT, DJL, CGP, PS, DF, and ZDN provided reagents and intellectual contribution. GR and MP wrote the paper with input from all authors.

## Data and materials availability

All accession codes, unique identifiers, and web links for publicly available datasets are available within the main text or the supplementary materials. Plasmids used in the study will be deposited to Addgene upon acceptance of the manuscript. All original data are available from the corresponding authors upon request. The raw mass spectrometry files are accessible at https://massive.ucsd.edu/ under the data set ID: MSV000091006 or at http://www.proteomexchange.org/ under data set ID: PXD039228 using the following credentials: Username: MSV000091006_reviewers, Password: Greg_turboID. The macro used for recruitment analysis for BER and MMR pathway components can be accessed at https://github.com/FenyoLab/recruitment_panel.

## Inclusion and diversity statement

We support inclusive, diverse, and equitable conduct of research.

## Supplementary Materials

Fig. S1 - Related to Fig. 1

Fig. S2 - Related to Fig. 2

Fig. S3 - Related to Fig. 3

Fig. S4 - Related to Fig. 4

Fig. S5 - Related to Fig. 5

Fig. S6 - Related to Fig. 6

Supplementary Figure Legends

Table S1 - Related to Fig. 2, TurboID-Cyclin D1 proteomics dataset

Table S2 - Related to Fig. 3, TurboID-PCNA proteomics dataset

## Supplementary Figure Legends

**Figure S1.**

A) Representative images of Fig. 1A. The green color shows the EGFP-tagged cyclins and red shows the mPlum-PCNA used to identify S phase cells and to highlight the sites of micro-irradiation.

B) U-2OS cells stably expressing mAzGreen-cyclin D1/D2 or D3 and mPlum-PCNA were pre-sensitized with BrdU for 24 hours and subjected to 405 nm laser induced damage. DNA damage recruitment dynamics were captured by live-cell imaging. S phase cells and the DNA damage site were identified based on the presence of PCNA foci. Relative fluorescence values and images were acquired every 5 seconds for 4 minutes. For each condition, ≥25 cells were evaluated from 3 independent experiments. Mean relative fluorescence values and standard errors were plotted against time. Representative images are shown next to the graphs. Times are indicated in seconds.

C) U-2OS cells stably expressing mAzGreen-cyclin D1 and mPlum-PCNA were pre-sensitized with BrdU for 24 hours and subjected to 405 nm laser induced damage using a confocal microscope. Cells were treated as in *(B)*. Representative images show recruitment to sites of laser-microirradiation at the indicated time in S and non-S phase cells.

D) The efficiency of the siRNA knock-downs is shown using immunoblotting for Fig. 1B.

E) RPE1 cells stably expressing mAzGreen-cyclin D1 and mPlum-PCNA were transfected with siRNAs targeting p21, p27, p57 or a non-targeting control (siNT). Cells were pre-sensitized with BrdU for 24 hours and subjected to 405 nm laser induced damage. Cells were treated and analyzed as in *(B)*. Representative images are shown below the graph. The efficiency of the siRNA knock-downs is shown using immunoblotting next to the graph.

F) Parental (*p21*^+/+^) or *p21*^-/-^ U-2OS cells stably expressing mAzGreen-cyclin D1 or NLS-mAzGreen-cyclin D1 and mPlum-PCNA were pre-sensitized with BrdU for 24 hours and subjected to 405 nm laser induced damage. Cells were treated and analyzed as in *(B)*. Representative images are shown next to the graphs.

G) U-2OS cells were transfected with the indicated FFSS-cyclin D1 constructs or an EV. Cell lysates were immunoprecipitated with an anti-FLAG resin, followed by elution using 3× FLAG peptide. Immunoprecipitates were probed with indicated antibodies.

H) Confocal images showing the localization pattern of the mAzGreen-cyclin D1 and the mAzGreen-cyclin D1(HP) construct with or without additional localization signals (NLS – nuclear localization signal, NES – nuclear export signal).

I) HEK 293T cells were co-transfected with the indicated FFSS-cyclin D1 constructs or an EV and either HA-ESα (Estrogen Receptor 1), used as a positive binding control, or HA-GR (Glucocorticoid Receptor), used as a negative binding control. Cell lysates were immunoprecipitated with an anti-FLAG resin, followed by elution using 3× FLAG peptide. Immunoprecipitates were probed with the indicated antibodies.

J) U-2OS cells stably expressing EGFP-cyclin D1 and mPlum-PCNA or EGFP-cyclin E1 and mPlum-PCNA were transfected with siRNAs p21 or a non-targeting control (siNT), pre-sensitized with BrdU for 24 hours and subsequently synchronized into G1. Cells were then subjected to 405 nm laser induced damage. Cells were treated and analyzed as in *(B)*.

K) U-2OS cells stably expressing mAzGreen-cyclin D1 and mPlum-PCNA were pre-sensitized with BrdU for 24 hours and subsequently synchronized into G1. Cells were then subjected to 405 nm laser induced damage with or without 2 hours of pre-treatment with palbociclib (CDK4/6i). Cells were treated and analyzed as in *(B)*. Representative images are shown next to the graphs.

L) Left: HEK 293T cells were co-transfected with FFSS-cyclin D1 or an EV and the indicated HA-tagged p21 constructs. p21(Δ17-21) denotes a mutant with a deletion within the indicated residues. p21(RxL) denotes mutations: R19A, R20A, L21A, F22A. p21(ΔPCNA) denotes mutations M147A, D149A, F150A. Cell lysates were immunoprecipitated with an anti-HA resin, followed by elution using HA peptide. Immunoprecipitates were probed with the indicated antibodies. Right: U-2OS cells were transfected with the indicated HA-tagged p21 constructs. Cell lysates were immunoprecipitated with an anti-HA resin, followed by elution using HA peptide. Immunoprecipitates were probed with the indicated antibodies.

M) HEK 293T cells were co-transfected with FFSS-cyclin D1 or EV and the indicated amount of HA-p21. Cell lysates were immunoprecipitated with an anti-FLAG resin, followed by elution using 3× FLAG peptide. Immunoprecipitates were probed with indicated antibodies.

N) U-2OS cells stably expressing p21-mCerulean constructs and mPlum-PCNA were pre-sensitized with BrdU for 24 hours and subjected to 405 nm laser induced damage using a confocal microscope. Cells were treated and analyzed as in *(B)*. Representative images are shown next to the graphs.

**Figure S2.**

A) G1-synchronized U-2OS and RPE1 cells were treated with the indicated DNA damaging agents. Cells were then fractionated into soluble and chromatin fractions, and lysates were immunoblotted as indicated. The figure shows the soluble fraction while Fig. 2A shows the chromatin fraction.

B) G1-synchronized parental (*p21*^+/+^) and *p21*^-/-^ U-2OS cells stably expressing mAzGreen-cyclin D1 or NLS-mAzGreen-cyclin D1 were treated with 100 μM H_2_O_2_ for 15 minutes. Cells were then fractionated into soluble and chromatin fractions and immunoblotted as indicated.

C) Parental (*p21*^+/+^) and *p21*^-/-^ U-2OS cells were transfected with siRNAs targeting p57 or a non-targeting control (NT), and subsequently synchronized in G1. Cells were then treated with 100 μM H_2_O_2_ for 15 minutes before fractionated into soluble and chromatin fractions, and lysates were immunoblotted as indicated.

D) 2-6-3 U-2OS cells were transfected with the indicated constructs and immunostained for γH2A.X (grey). White arrowheads indicate the position of the LacO in the nucleus (outlined by white dashed line). Quantification of the γH2A.X relative mean fluorescence intensity (RMFI) was carried out from 3 independent experiments as in ^83^ and visualized on violin plots. Dotted lines represent upper and lower quartiles while dashed lines represent the median on the plots. Scale bar represents 2 μm.

E) 2-6-3 U-2OS cells were transfected with the indicated constructs and immunostained for OGG1 (grey). White arrowheads indicate the position of the LacO in the nucleus (outlined by white dashed line). Quantification of the OGG1 relative mean fluorescence intensity (RMFI) was carried out from 3 independent experiments and visualized on violin plots. Dotted lines represent upper and lower quartiles while dashed lines represent the median on the plots. Scale bar represents 2 μm.

F) 2-6-3 U-2OS cells were transfected with the indicated constructs and immunostained for γH2A.X (green) and cyclin D1 (grey). The latter was detected using the DCS6 anti-cyclin D1 mouse monoclonal antibody. White arrowheads indicate the position of the LacO in the nucleus (outlined by white dashed line). Quantification of the cyclin D1 relative mean fluorescence intensity (RMFI) was carried out from 3 independent experiments and visualized on violin plots. Dotted lines represent upper and lower quartiles while dashed lines represent the median on the plots. Scale bar represents 2 μm.

G) 2-6-3 U-2OS cells were transfected with the indicated constructs and with siRNAs targeting p21 or a non-targeting control (NT), immunostained for γH2A.X (green) and cyclin D1 (grey). Latter was detected using the DCS6 mouse monoclonal antibody. Quantification of cyclin D1 relative mean fluorescence intensity (RMFI) was carried out from 3 independent experiments and visualized on violin plots. Dotted lines represent upper and lower quartiles while dashed lines represent the median on the plots. Scale bar represents 2 μm.

H) Immunoblot image related to Fig. 2H. U-2OS cells stably expressing HA-TurboID, HA-TurboID-cyclin D1 (5 nm linker) or HA-TurboID-13x-cyclin D1 (25 nm linker) were treated with biotin and H_2_O_2_ for 1.5 hours as indicated. Cells were then fractionated into soluble and chromatin fractions, and lysates were immunoblotted as indicated.

I) Schematics showing how the HA-TurboID-cyclin D1 constructs were used to biotinylate proteins in its proximity upon oxidative DNA damage (represented by a base colored in red) compared to HA-TurboID alone.

**Figure S3.**

A) Immunoblot related to Fig. 3A. RPE1 cells stably expressing NLS-HA-TurboID or NLS-HA-TurboID-PCNA were transfected with siRNAs targeting D-type cyclins or a non-targeting control (siNT), and subsequently synchronized in G1. Cells were treated with biotin and H_2_O_2_ for 1.5 hours as indicated. Cells were then fractionated into soluble and chromatin fractions, and lysates were immunoblotted as indicated. This figure shows the expression levels of the various constructs and the recruitment of TurboID-PCNA to the chromatin upon oxidative damage.

B) Left: Schematics showing how the NLS-HA-TurboID-PCNA construct was used to biotinylate proteins in its proximity upon oxidative DNA damage (represented by a base colored in red) in the presence or absence of D-type cyclins. Right: Volcano plot of protein change comparing NLS-HA-TurboID and NLS-HA-TurboID-PCNA. Known PCNA interactors (highlighted in red) are significantly enriched in the PCNA samples (p = 2.42e-06, Fisher’s exact test). Proteins were selected based on a fold change greater than or equal to 2 and an (uncorrected) *p*-value less than or equal to 0.05. Raw values are found in Supplementary Table S2.

C) U-2OS cells were transfected with either non-targeting (siNT) or siRNA targeting D-type cyclins or KIPs, and subsequently synchronized in G1. Cells were then treated with 100 μM H_2_O_2_ before fractionating into soluble and chromatin fractions. PCNA was immunoprecipitated from the chromatin fraction and co-purified proteins were immunoblotted as indicated (left panel). Right panels show chromatin and soluble fractions.

D) G1-synchronized parental (*AMBRA1*^+/+^) or *AMBRA1*^-/-^ U-2OS cells were treated with 100 μM H_2_O_2_ before fractionating into soluble and chromatin fractions. PCNA was immunoprecipitated from the chromatin fraction and co-purified proteins were immunoblotted as indicated (left panel). Right panels show chromatin and soluble fractions.

E) U-2OS cells stably expressing mCherry-tagged BER pathway components or AcGFP-tagged MMR pathway components were transfected with either non-targeting or siRNA targeting D-type cyclins, and subsequently synchronized in G1. DNA damage recruitment dynamics were captured by live-cell imaging. Relative fluorescence values and images were acquired every 5 seconds for 20 minutes. Mean relative fluorescence values and standard errors were plotted against time. Times are indicated in seconds.

**Figure S4.**

A) Representative images of the comet assay shown in Fig. 4A,B.

B) RPE1 cells were transfected with either siRNAs targeting D-type cyclins, KIPs, or a non-targeting siRNA control (siNT), and subsequently synchronized them in G1. Cells were then treated with H_2_O_2_ for 15 minutes or left untreated (UT). Samples treated with H_2_O_2_ were allowed to recover in media without H_2_O_2_ for the indicated time. Lysates were blotted with the indicated antibodies.

C) RPE1 cells were transfected with either siRNAs targeting the indicated proteins or a non-targeting siRNA control (siNT), and subsequently synchronized in G1. Where indicated, cells were treated with H_2_O_2_ for 30 minutes and then allowed to recover for 1.5 hours in DMEM supplemented with EdU. The EdU signal was normalized to the untreated siNT samples. Measurements were carried out from 3 independent experiments and visualized on violin plots. Dashed lines represent the median on the plots. The efficiency of the siRNA knockdowns is shown using immunoblotting on the right.

D) Schematics of the transcriptional mutagenesis-based fluorescent MMR reporters [G:G and A:C] are shown on the left. The base pairs shown correspond to sites that code for a key amino acid of the chromophores of the fluorescent proteins. The transcribed strand is on the top.

E) TK6 MLH1^+/+^ or TK6 MLH1^-/-^ cells nucleoporated with the indicated FM-HCR reporter plasmids. 20 hours later, the cells were subjected to FACS analysis. Repair capacity for each sample was normalized to MLH1^+/+^ and is represented as normalized repair capacity. Graphs show average and standard deviation from at least 3 independent experiments.

F) RPE1 cells were transfected with siRNAs targeting MSH2 and MLH1 or a non-targeting control (siNT) and serum-starved to arrest them in G0. The cells were then nucleoporated with the indicated FM-HCR reporter plasmids and released into serum containing media. Ten hours later, cells, now in G1, were subjected to FACS analysis. Repair capacity for each sample was normalized to siNT and is represented as relative repair capacity. Graphs show average and standard deviation from at least 3 independent experiments. The efficiency of the siRNA knock-downs is shown using immunoblotting on the right.

G) Stably transduced RPE1 cells, harboring a doxycycline inducible FLAG-cyclin D1/STREP-cyclin D2/HA-cyclin D3 cassette or an HA-p21, HA-p21(ΔPCNA), EV constructs, were serum-starved to arrest them in G0 while the transgenes were induced by doxycycline. The cells were then nucleoporated with the indicated FM-HCR reporter plasmids and released into serum containing media. Ten hours later, cells, now in G1, were subjected to FACS analysis. Repair capacity for each sample was normalized to EV and is represented as normalized repair capacity. Graphs show average and standard deviation from at least 3 independent experiments. The efficiency of the doxycycline induction for each gene is shown using immunoblotting on the right.

H) Parental (*AMBRA1*^+/+^) or *AMBRA1*^-/-^ RPE1 cells (clones #KO1 and #KO2) were transfected with siRNA targeting KIPs or a non-targeting control (siNT) and serum-starved to arrest them in G0. Cell were then nucleoporated with the indicated FM-HCR reporter plasmids and released into serum containing media. Ten hours later, cells, now in G1, were subjected to FACS analysis. Repair capacity for each sample was normalized to parental (*AMBRA1*^+/+^) cells transfected with siNT and is represented as normalized repair capacity. Graphs show average and standard deviation from at least 3 independent experiments. The efficiency of the siRNA knock-downs is shown using immunoblotting on the right.

I) Representative FACS gating strategy used for the FM-HCS assays. Gating for live cells (a) and single cells (b) are shown on the left. Upper panels show non-transfected (c), EGFP (d), mPlum (e), and EGFP plus mPlum (f) transfected cells in the EGFP / mPlum channels. The lower panel shows non-transfected (g), mOrange (h), EBFP (i), and mOrange plus EBFP (j) transfected cells in the mOrange / EBFP channels.

**Figure S5.**

A) RPE1 cells were transfected with either non-targeting or siRNA targeting CDK4 and 6, and subsequently synchronized into G1. Where indicated, cells were pre-treated with palbociclib (CDK4/6i) for 2 hours before treatment with 100 μM H_2_O_2_ and fractionation into soluble and chromatin fractions. PCNA was immunoprecipitated from the chromatin fraction and co-purified proteins were immunoblotted as indicated (left panel). Right panels show chromatin and soluble fractions.

B) The graph shows the results DNA repair synthesis assay. RPE1 cells were transfected with siRNAs targeting CDK4 and 6 or a non-targeting control (siNT), and subsequently synchronized into G1. Where indicated, cells were then pre-treated with palbociclib (CDK4/6i) for 2 hours, before treating them with H_2_O_2_ for 30 minutes. After 1.5 hours of recovery in media with EdU (with or without palbociclib), cells were processed for the assay. The EdU signal were normalized to the siNT samples. Measurements were carried out from 3 independent experiments and visualized on violin plots. Dashed lines represent the median on the plots. The efficiency of the siRNA knock-downs and the palbociclib (CDK4/6i) treatment are shown using immunoblotting on the right.

C) Parental (*AMBRA1*^+/+^) or *AMBRA1*^-/-^ RPE1 cells (clones #KO1 and #KO2) and serum-starved to arrest them in G0. Where indicated, cells pre-treated with palbociclib (CDK4/6i) for 1 hour before the cells were nucleoporated with the indicated FM-HCR reporter plasmids and released into serum containing media in the presence or absence of palbociclib. Ten hours later, cells, now in G1, were subjected to FACS analysis. Repair capacity for each sample was normalized to parental (*AMBRA1*^+/+^) cells and is represented as normalized repair capacity. Graphs show average and standard deviation from at least 3 independent experiments.

D) RPE1 or U-2OS cells were transfected with siRNAs targeting D-type cyclins, p21 or a non-targeting control (siNT), and subsequently synchronized in G1. Graph shows relative mean p21 mRNA levels of these samples as measured by qPCR from 3 independent measurements. Error bars represent standard deviation. The efficiency of the siRNA knock-down of the D-type cyclins and their effect on p21 protein levels are shown using immunoblotting below.

E) Parental (*AMBRA1*^+/+^) or *AMBRA1*^-/-^ U-2OS cells (clones #KO1 and #KO2) were treated with cycloheximide (CHX) and MG132 as indicated, and lysates were blotted with the indicated antibodies. The graph shows the quantification of p21 levels from three independent experiments. Error bars indicate standard error.

F) *AMBRA1*^-/-^ U-2OS cells (clones #KO1 and #KO2) were transfected with siRNAs targeting D-type cyclins, or a non-targeting control (siNT) before treated with cycloheximide (CHX) and MG132 as indicated. Lysates were blotted with the indicated antibodies. The graph shows the quantification of p21 levels from three independent experiments. Error bars indicate standard error.

G) U-2OS cells were transfected with siRNAs targeting D-type cyclins, or a non-targeting control (siNT), and subsequently synchronized in G1. Cells were then treated with cycloheximide (CHX) and MG132 or MLN4924 as indicated, and lysates were blotted with the indicated antibodies. The graph shows the quantification of p21 levels from three independent experiments. Error bars indicate standard error.

H) Parental (WT) or *CCND1/D2/D3* knockout MEF cells were treated with cycloheximide (CHX) and MG132 as indicated, and lysates were blotted with the indicated antibodies. The graph shows the quantification of p21 levels from three independent experiments. Error bars indicate standard error.

I) Stably transduced U-2OS cells, harboring a doxycycline inducible HA-p21, HA-p21(ΔPCNA) or an EV, were synchronized into G1-phase during which the transgenes were induced by doxycycline. G1-synchronized cells were then treated with 100 μM H_2_O_2_ before fractionating into soluble and chromatin fractions. PCNA was immunoprecipitated from the chromatin fraction and co-purified proteins were immunoblotted as indicated. Right panels show chromatin and soluble fractions.

**Figure S6.**

A) Cell synchronization. Left: Immunoblot shows cell cycle markers in parental (*AMBRA1*^+/+^) and *AMBRA1*^-/-^ U-2OS cell lysates at the indicated time points after releasing them from a 14-hour nocodazole block using mitotic shake-off. Cell cycle stage of each lane is indicated at the bottom. Right: Immunoblot shows cell cycle markers in parental (*AMBRA1*^+/+^) and *AMBRA1*^-/-^ RPE1 lysates at the indicated time points after releasing them from a 72 hours-long serum starvation. Cell cycle stage of each lane is indicated at the bottom.

B) S phase-synchronized parental (*AMBRA1*^+/+^) or *AMBRA1*^-/-^ U-2OS cells (clones #KO1 and #KO2) were fractionated into soluble and chromatin fractions. PCNA was immunoprecipitated from the chromatin fraction and co-purified proteins were immunoblotted as indicated (left panel). Right panels show chromatin and soluble fractions. Asterix denoted a non-specific band.

C) S phase-synchronized parental (*AMBRA1*^+/+^) or *AMBRA1*^-/-^ U-2OS cells (clones #KO1 and #KO2) were treated with 100 μM H_2_O_2_ before fractionated into soluble and chromatin fractions. PCNA was immunoprecipitated from the chromatin fraction and co-purified proteins were immunoblotted as indicated (left panel). Right panels show chromatin and soluble fractions.

D) Representative FACS gating strategy used for the mCherryOFF assay. Gating for live cells (a) and single cells (b) are shown on the top. Lower panels show non-transfected cells (c), EGFP (d), mCherry (e), EGFP plus mCherry (f) and mCherryOFF reporter expressing cells (g) in the EGFP / mCherry channels.

E) Graphs show NanoLuc signal in U-2OS cells, normalized to cell viability. NanoLuc signal was evaluated 96 hours after transfecting cells with siRNAs targeting MSH2 along with MLH1 or a non-targeting control (siNT). Graphs show average and standard deviation from at least 3 independent experiments. Efficiency of the siRNA silencing is shown using immunoblotting next to the graphs.

F) Graphs show NanoLuc signal in MCF10Am cells, normalized to cell viability from 3 independent experiments. Cells were treated and analyzed as in *(D)*. Efficiency of the siRNA silencing is shown using immunoblotting next to the graphs.

