## Supplementary figures and images for "D-type cyclins regulate DNA mismatch repair in the G1 and S phases of the cell cycle, maintaining genome stability"

### Supplementary Figure 1

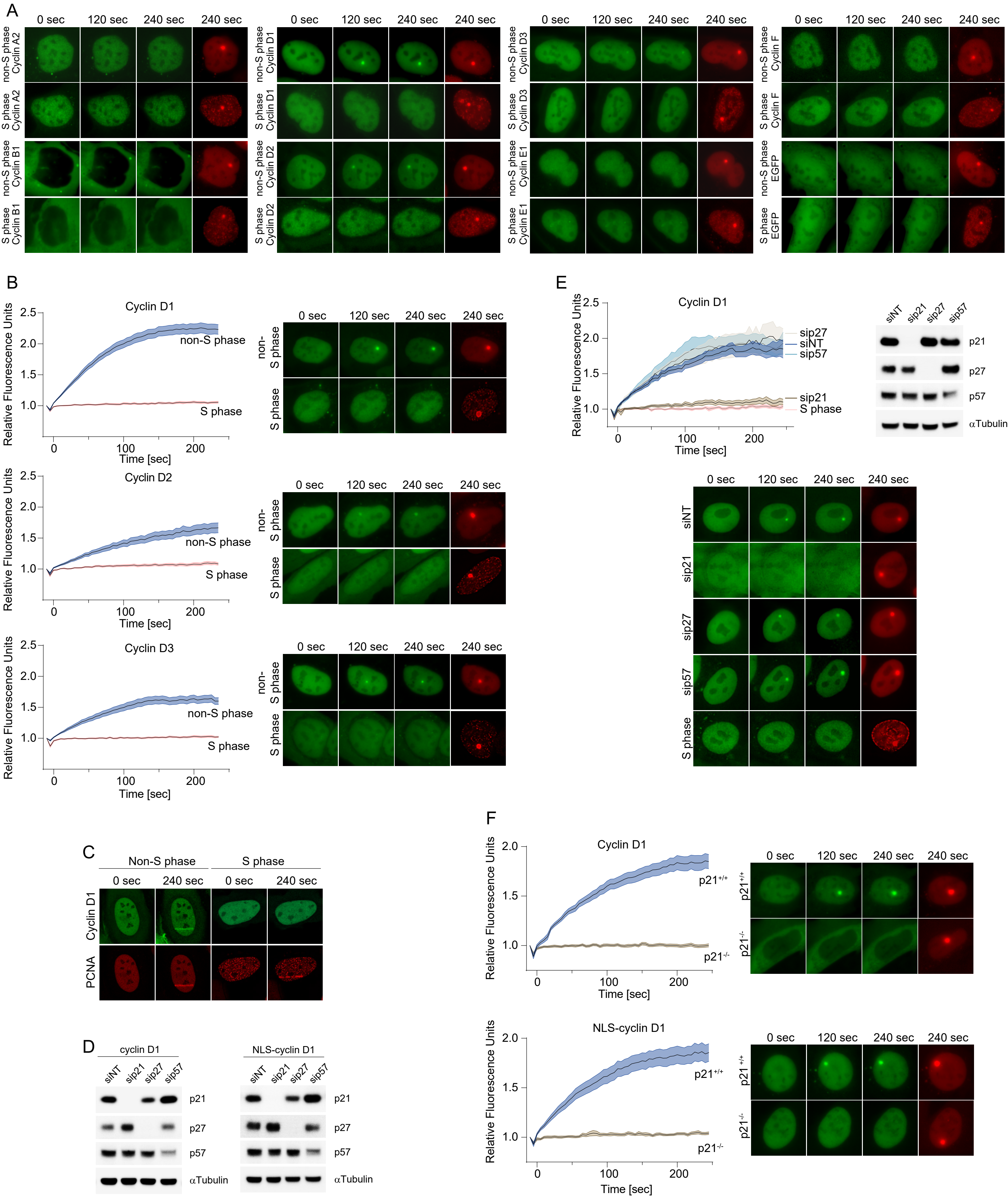

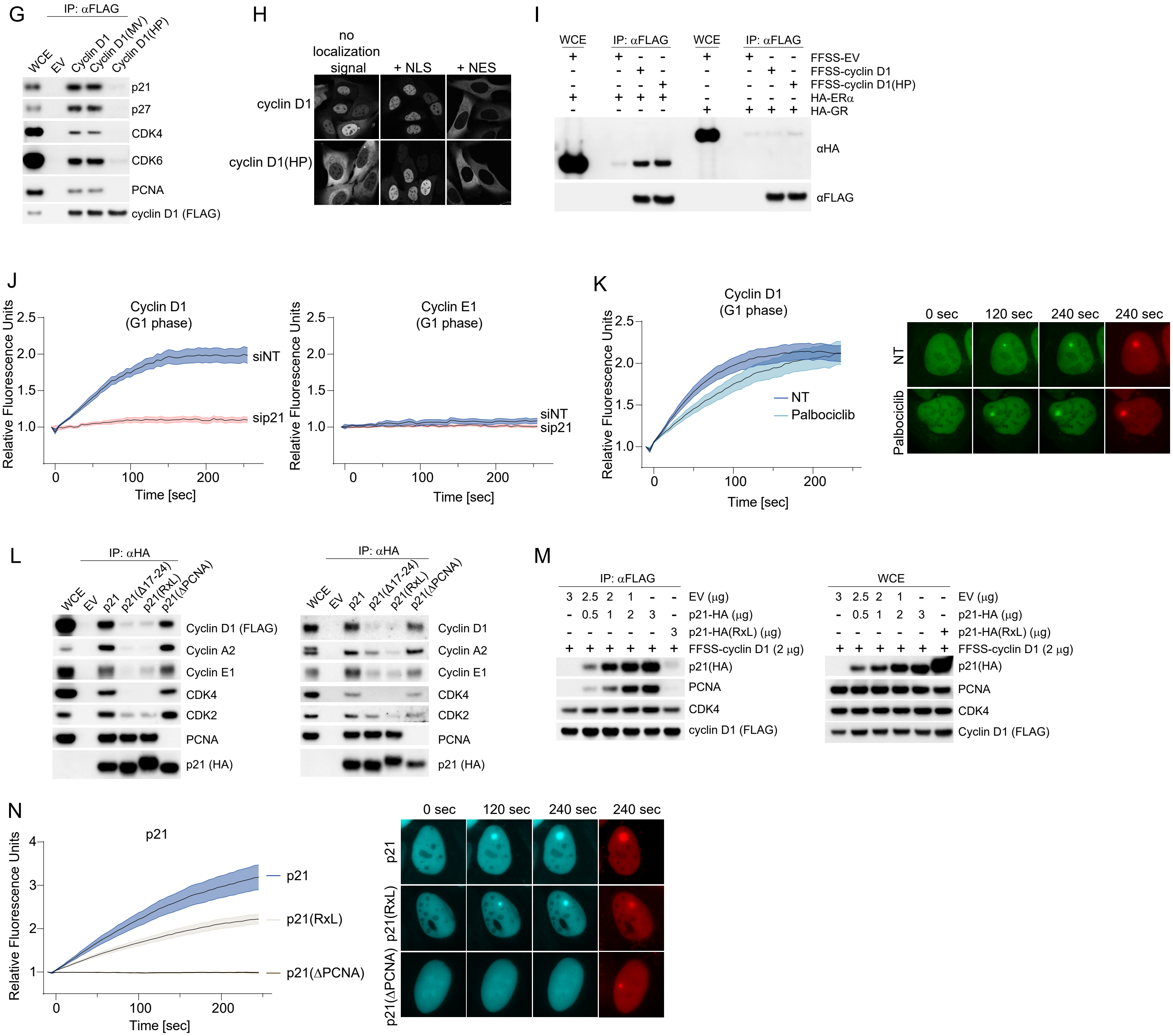

### Supplementary Figure 2

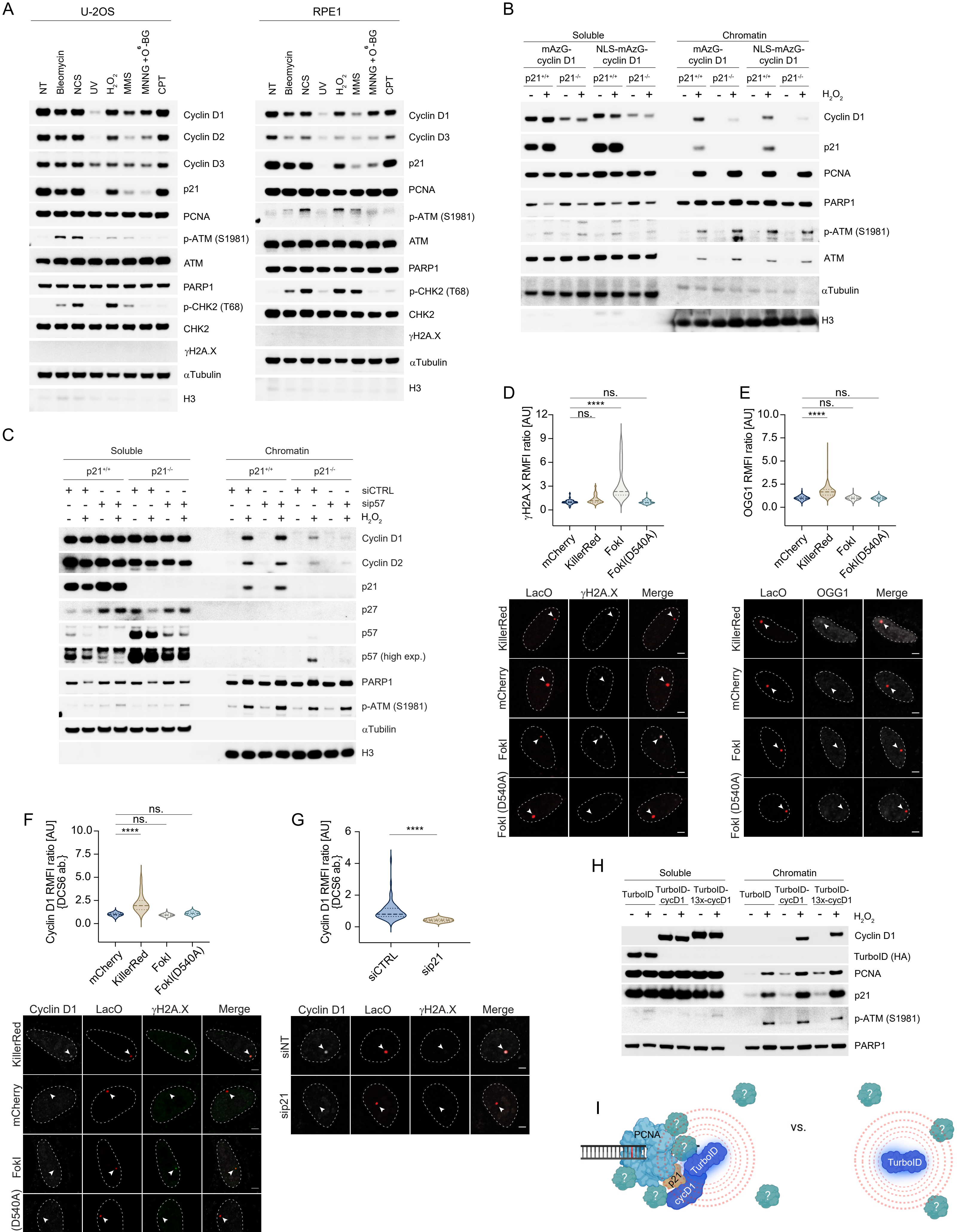

### Supplementary Figure 3

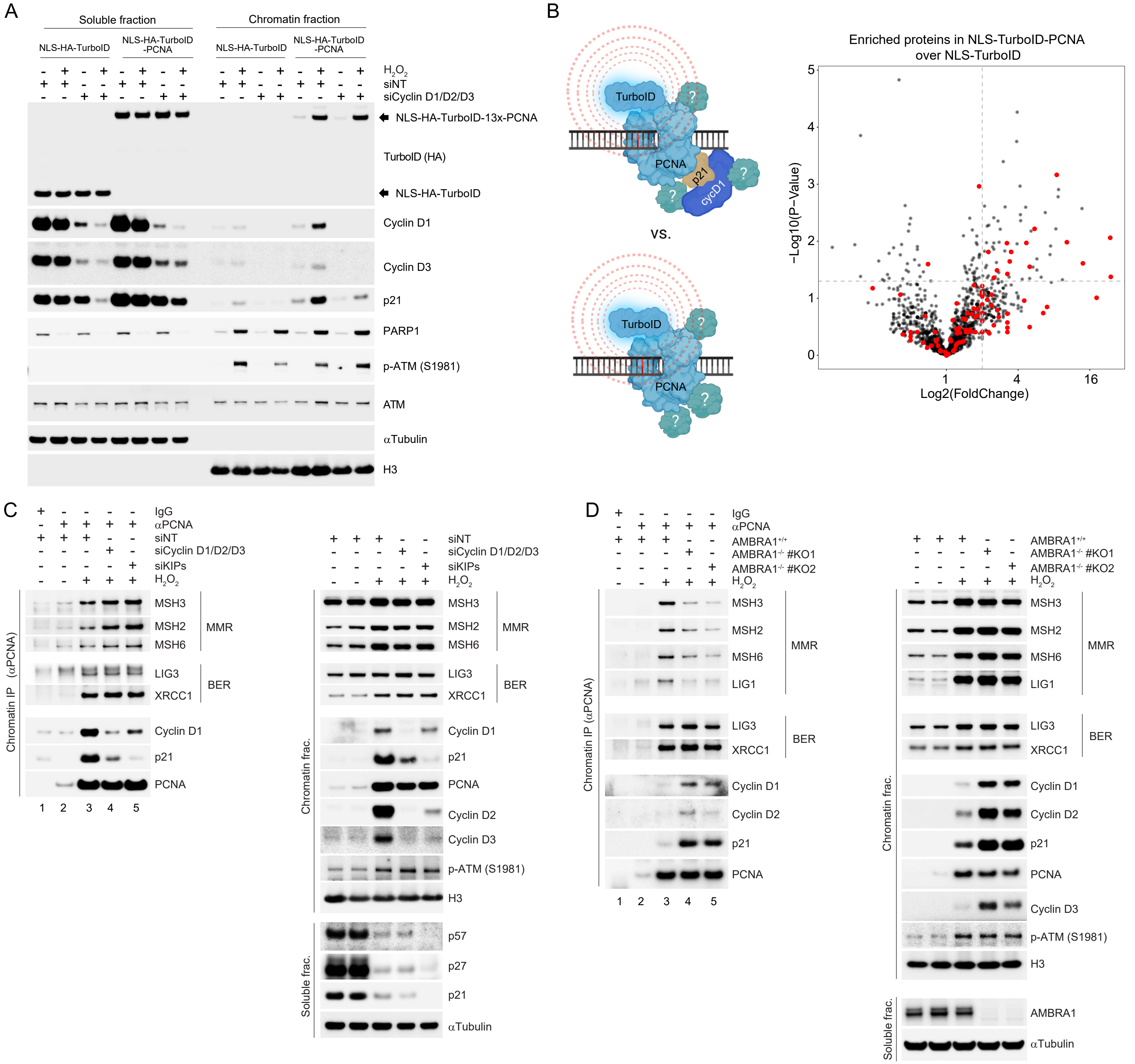

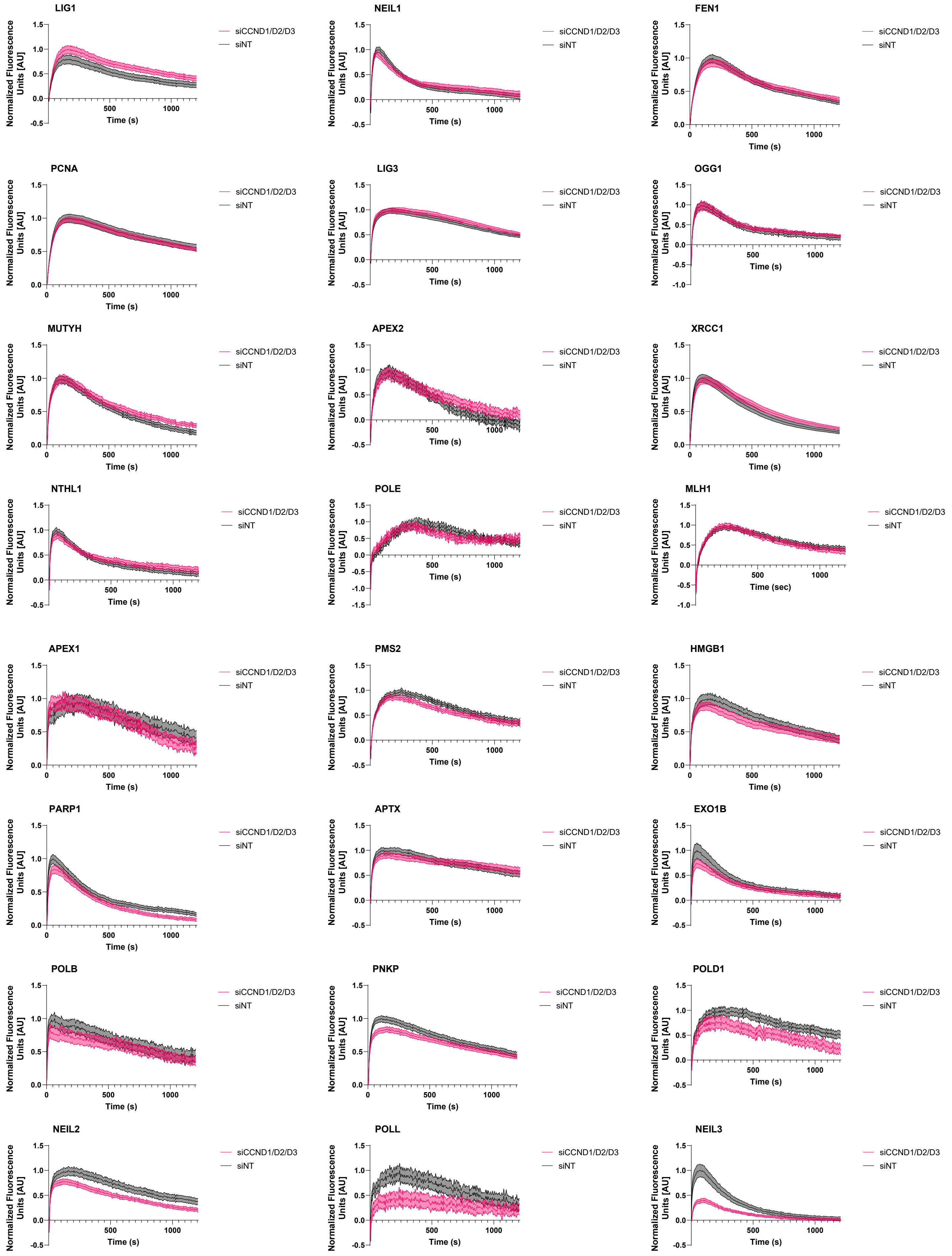

### Supplementary Figure 4

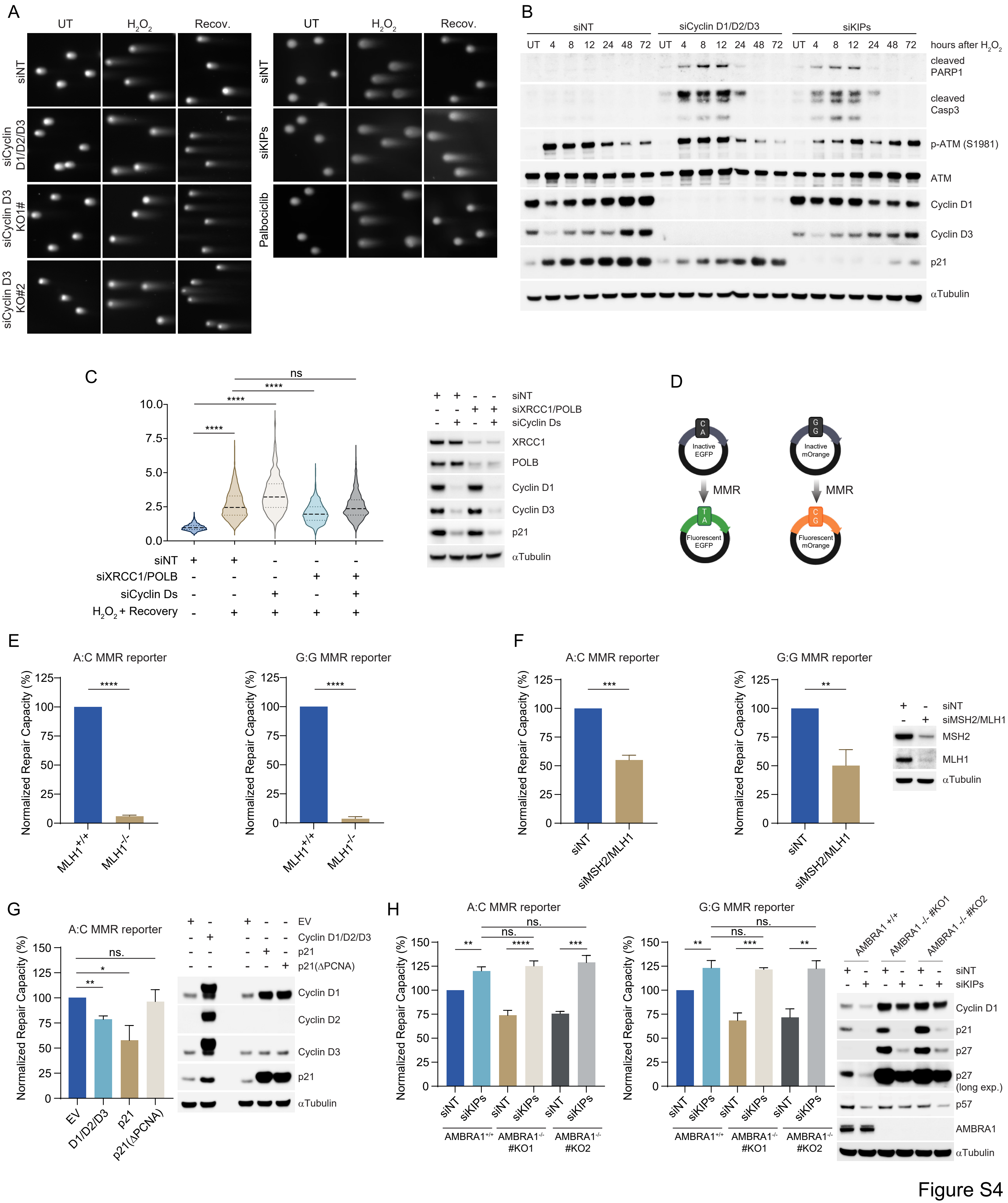

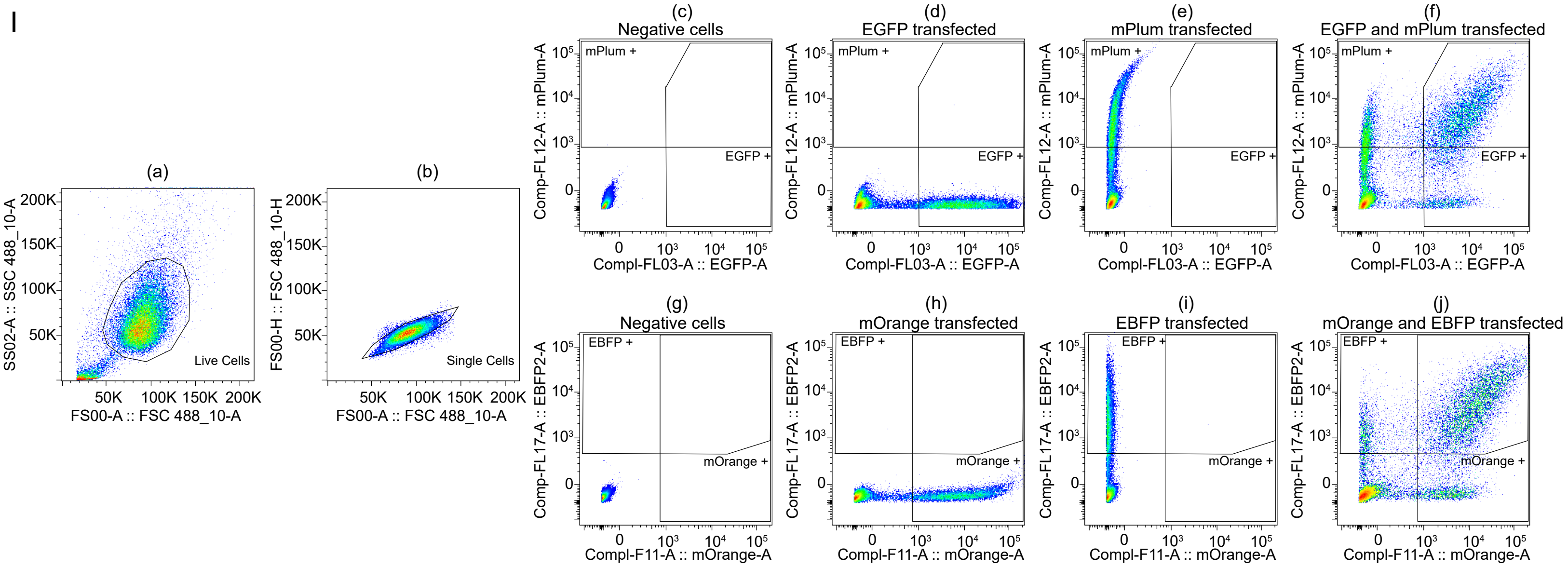

### Supplementary Figure 5

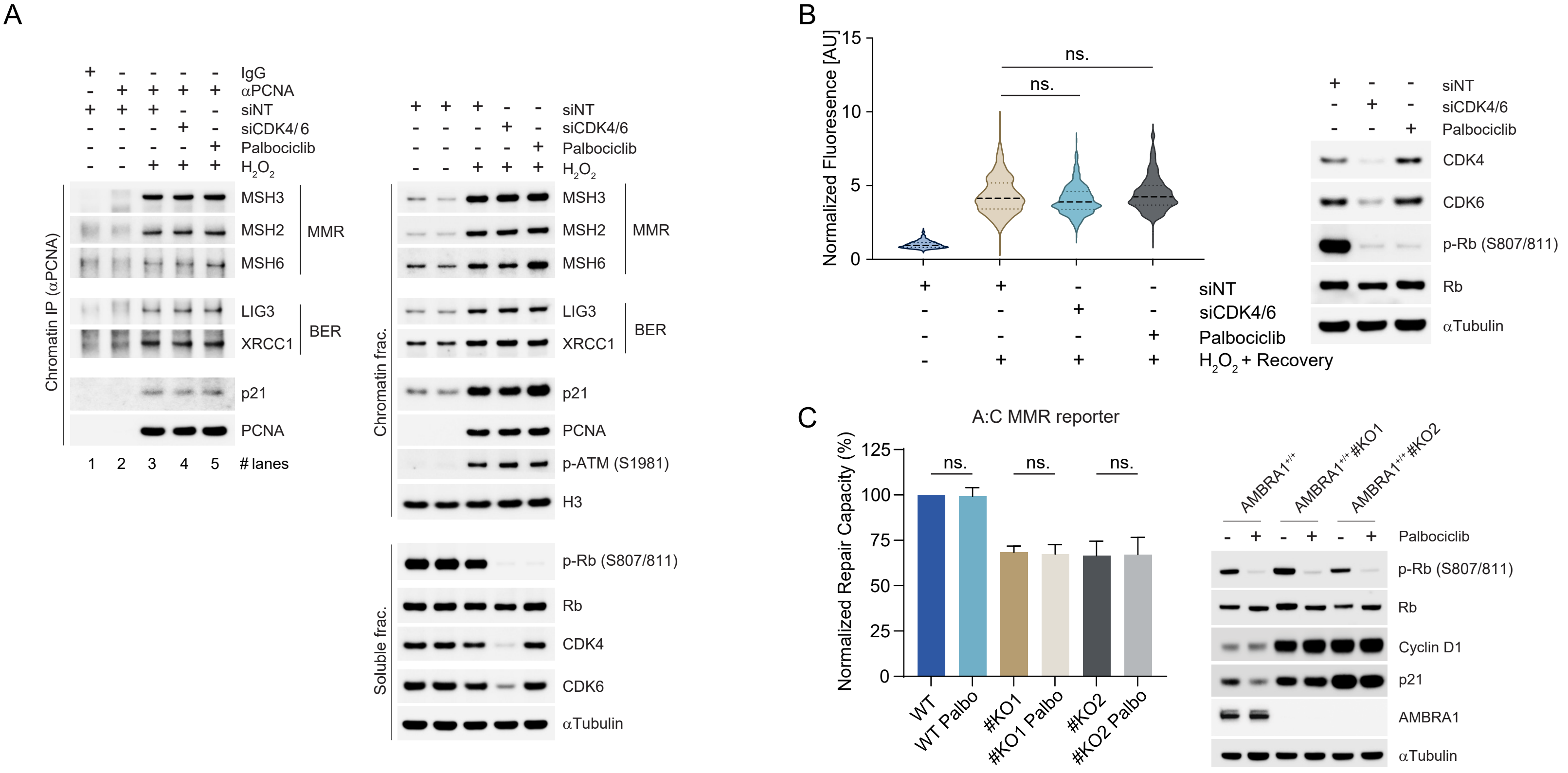

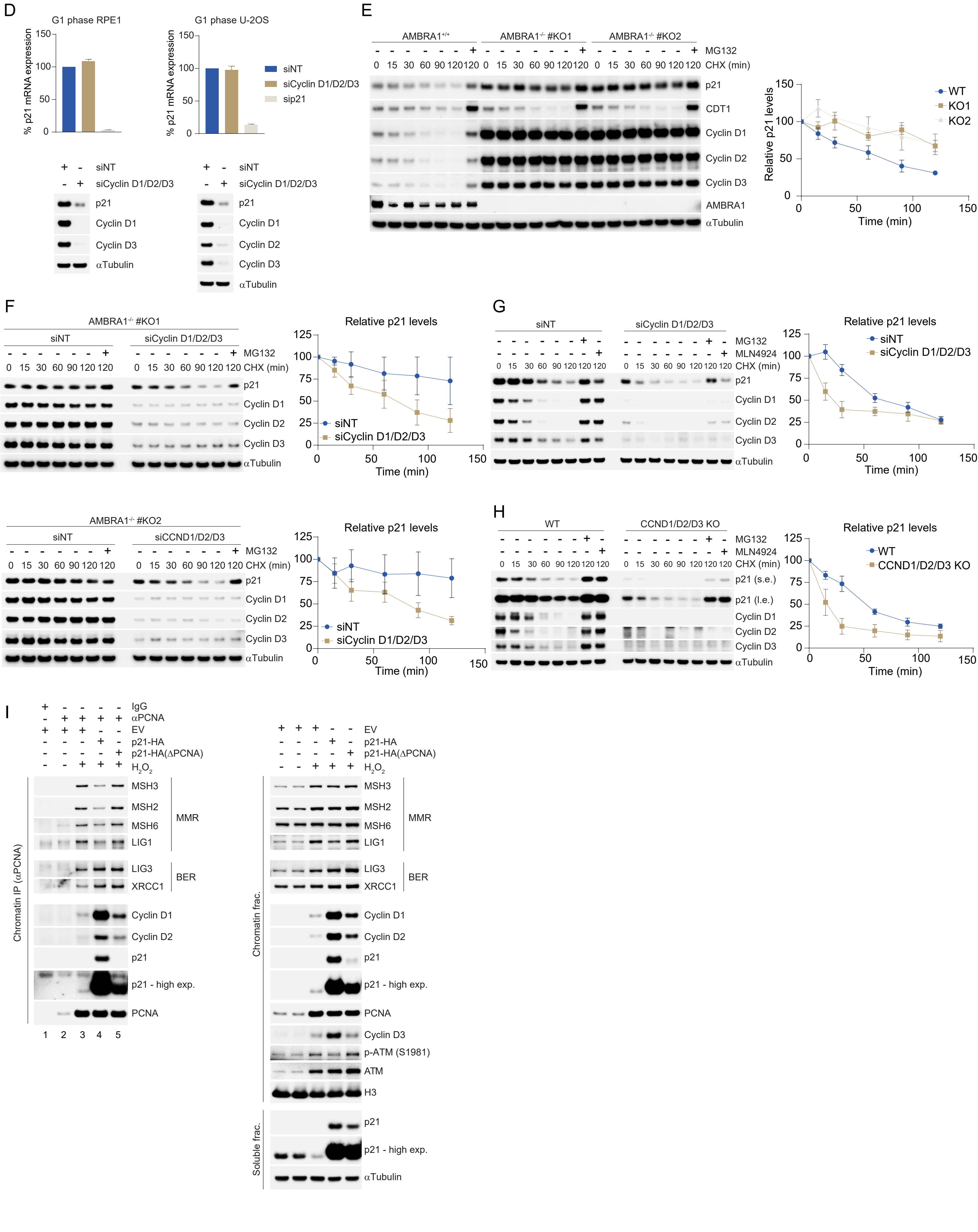

### Supplementary Figure 6

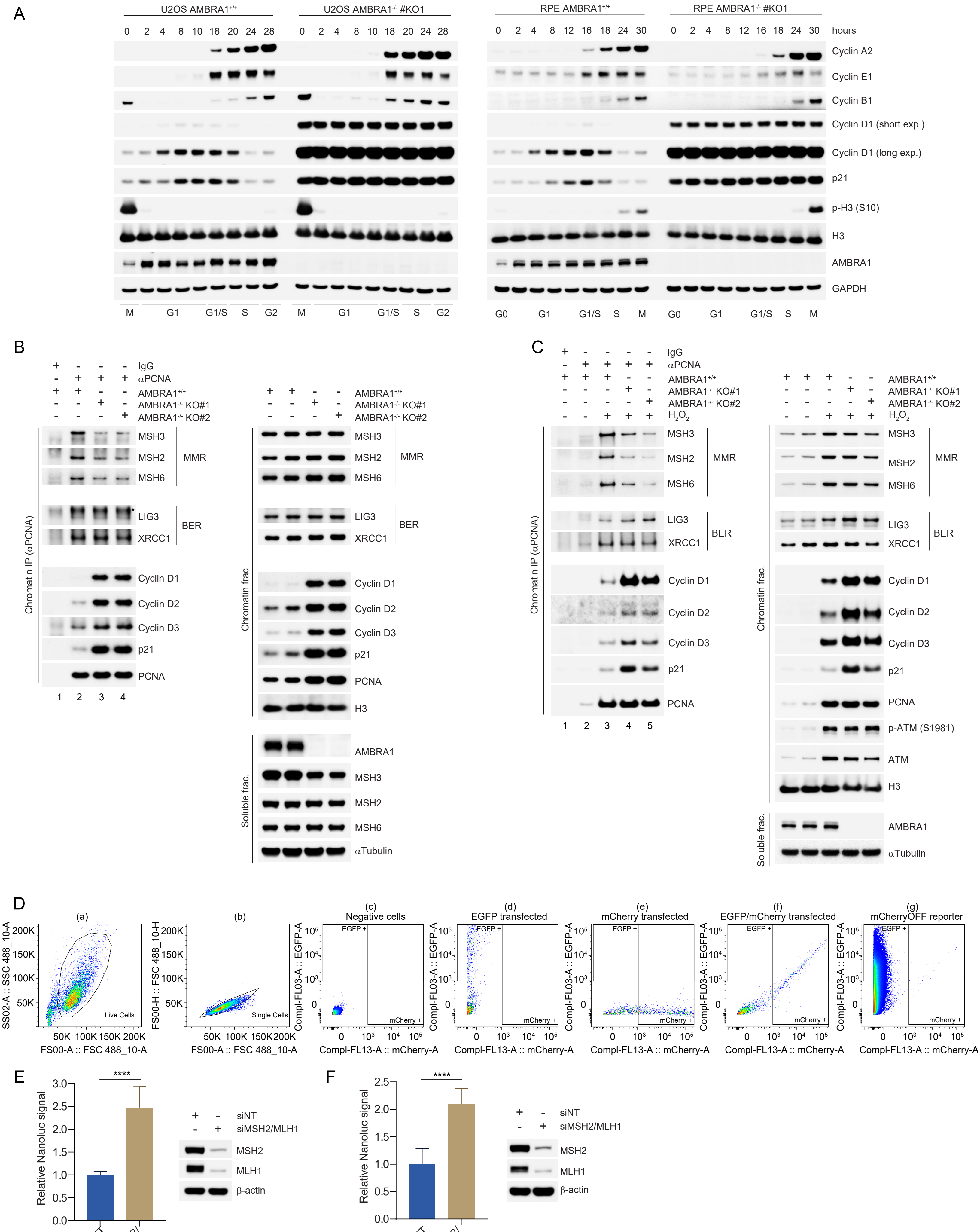
